# Sleep Deprivation in Mice: Looking Beyond the Slow Wave Rebound

**DOI:** 10.1101/2024.10.31.621300

**Authors:** Tárek Zoltán Magyar, Orsolya Szalárdy, Róbert Bódizs

## Abstract

Sleep is a fundamental process by which the brain achieves an optimal computational regime; it has been suggested that criticality is an appropriate theoretical framework in which such a process can be understood. Studying the critical and fractal dynamics of the brain involves modelling the nonlinearity of brain signals (such as EEG or ECoG) yielding, among others, the following metrics: spectral slope, spectral intercept, spectral knee, and normalized spectral entropy. Therefore, the present study investigates the nonlinear and critical dynamics of the brain in relation to sleep deprivation in mice, by comparing the sensitivity of the above-mentioned metrics to classical band-limited spectral indices. Mice were exposed to a 9 day-long sleep deprivation paradigm with baseline, sleep deprivation, and recovery phases. Spectral parameters were computed using the FOOOF algorithm. The results suggest that the classical approach (slow wave activity; 0.75-4.5 Hz) to neural signal processing differentiates between baseline sleep and rebound sleep only during the NREM phase. In contrast, the spectral slope and the spectral intercept both capture sleep deprivation related effects during REM and NREM episodes as well. This is particularly notable considering that the spectral knee is shifted towards higher frequencies, essentially rendering the spectral slope unreflective of slow wave activity – traditionally considered the biomarker of sleep homeostasis.

Lastly, normalized spectral entropy fails to differentiate between baseline sleep and sleep following sleep deprivation in mice. These results support the sensitivity of fractal spectral parameters indexing the intricate balance between sleep and wake states.

## Introduction

Earlier theoretical modelling and recent experimental evidence suggests that the primary function of sleep is the maintenance of criticality, that is the optimal computational regime within the central nervous system (1,2). The critical brain hypothesis assumes that brain dynamics may be characterised by a critical point interspersed between two fundamental macro states, a supra- and a suboptimal state (3–6). Criticality is a concept of statistical physics; it denotes a point at which a dynamical system behaves qualitatively differently, corresponding to a phase transition between states. That is to say, the hypothesis proposes that neural networks, and thus various facets of brain activity self-organize into and operate near a phase transition, at which information processing, adaptability, and computing properties are optimized (7). In practice, criticality can be understood as a boundary of neural parameter space separating two states of neural activity expressing qualitatively different behaviours. Alternatively, criticality describes “a border that separates networks with persistent but controlled activity from networks with pathological behaviour” (1). Scale invariant phenomena, such as power laws and fractals, are characteristic features of phase transitions (8). Furthermore, the collective behaviour of neurons can be described by avalanches of activity with features following power law distributions (3). The exponent of a power law (also known as slope on the double logarithmic plane) describes the functional relationship between variables as being relatively proportional irrespective of the initial sizes of the quantities involved, and it is essentially the manifestation of fractal-like properties (9,10). Consequently, assessing the power-law scaling, and thus fractality of a signal may give insight in terms of critical dynamics.

The two-process model, as its name suggests, postulates two processes that underlie sleep regulation: The C-process refers to the circadian rhythm and the S-process is the homeostatic pressure (11). A biomarker which is assumed to reflect on sleep homeostasis should implicitly also display adjustments related to sleep deprivation. Accordingly, a well-established biomarker of homeostatic sleep regulation during sleep recovery consecutive to prolonged wakefulness is the increase in the surface neurophysiological spectral power within the low or delta frequency range during sleep (11–13). Slow wave activity (SWA; 0.75-4.5 Hz, to some degree interchangeable with delta waves: 1-4 Hz) peaks during the deepest sleeping stages of the non-rapid eye movement (NREM) phase, and it is speculated to be related to sleep maintenance and to the restorative nature of sleep. Sleep deprivation has been associated to impaired frontoparietal functioning (14); and in accordance with that, it has been suggested that sleep EEG delta waves are imperative in the restoration of frontal and medial temporal regions (15). Therefore, the sleep deprivation-related enhancement of the SWA frequency range can be considered a further increase in sleep depth and a compensatory mechanism for the inappropriately prolonged wakefulness. However, classical methods of neurophysiological signal processing suffer from non-ideal statistical properties, namely, the lack of a reference point rendering inter-individual comparisons problematic (16). A proposed solution is based on the idea that surface neurophysiological signals demonstrate a fractal-like nature, and as a consequence, they can be decomposed into an aperiodic and possibly multiple periodic components on the frequency domain (16,17). This data-driven approach to signal processing yields measures of the power spectrum which are associated to age-effects and sex-differences (16), arousal level (18), task demand and brain state during sleep and wakefulness (19), sleep intensity (20) sleep staging (21,22), and homeostatic sleep regulation (23). As a general rule, a steeper EEG spectral slope indicates more intense sleep as inferred from the findings that factors and conditions known to associate with high SWA, like young age, deeper stages derived from rule-based scoring, early sleep cycles, and anterior recording sites are reliably reflected in the accelerated rate of decay of spectral power along the frequency axis (23).

The theory of Pearlmutter and Houghton (1) postulates that changes in synaptic efficacies throughout wakefulness alters the activity persistence and response rapidity of neural networks, shifting the brain to operate away from criticality. However, without exerting a force against such shifts, uncontrolled feedback, and thus anti-persistent behaviour, that is runaway oscillation into a supercritical state is at risk. Accordingly, sleep serves as a mechanism to keep neural networks operating near the critical boundary. That is to say, neural parameters vary within a dynamical range defined by a margin of safety, within which neural computations are optimized. As a response to sleep deprivation, however, the critical boundary shifts towards supercritical behaviour, resulting in an increased margin of safety at the expense of processing efficiency (giving rise to tiredness). In other words, the model predicts that wakefulness, as a function of time, progressively disrupts signatures of criticality towards a supercritical state; it has been shown that the brain during sustained wakefulness is indeed characterized by increased supercriticality (24,25). Therefore, it is safe to assume that as a recovery measure, sleep followed by prolonged wakefulness should be characterized by markers of enhanced compensation. Recent experimental evidence based on neuronal spiking measurements of rats cohere with this notion and suggests that waking experience progressively disrupts criticality and that sleep functions to restore critical dynamics (2). Moreover, the degree of deviation from criticality predicts future sleep/wake behaviour more accurately than SWA/delta activity (2). Indeed, the effect of extended wakefulness was mainly or exclusively tested on the grounds of the classical, band-limited spectral measures of EEG/cortical LFP, whereas the fractal spectral indices known to be closely related to the critical dynamics, were not tested.

The study of complex systems must be based on metrics that are intrinsically sensitive to nonlinear behaviour, such as informational entropy. Traditionally, high entropy signifies a system with a lot of randomness and uncertainty, while low entropy indicates a more predictable, ordered state.

However, a more recent school of thought (26,27) argues that a maximally random system (described by a white-noise function on the frequency domain) should be characterized by low values of entropy. Accordingly, the brain’s informational entropy is hypothesized to be the highest when the system is at a critical point (28–30). The brain is poised to process a wide range of inputs and adapt to unknown situations, increasing variability in neural activity. However, this variability is not chaotic; it is constrained by the need to maintain functional order and coordination. At criticality, the brain exhibits a high variability while remaining functionally coherent (not overly chaotic), at which state the brain optimizes information flow, allowing for efficient coding of information with minimal redundancy. Thus, informational entropy is maximized at the edge of chaos - neither too ordered nor too disordered. This concept provides an additional framework for neuroscientific investigation emphasizing the relevance of signal irregularity, which therefore assumes the brain to produce complex, nonlinear signals (31–33); and in accordance with that, various indices of entropy may be appropriate to characterize the seemingly hidden dynamics embedded in neural signals (34). Consonantly, neural complexity, as computed by various entropy metrics, has been associated to, including, but not limited to ADHD (35), autism (36), human intelligence (37) and aging (38). Finally, it has also been related to sleep onset latency (39), sleep quality (40), sleep spindle detection (41), used as a feature in sleep staging (42,43), and lastly revealed to reflect the balance between sleep- and alertness-promoting processes (44). Particularly interesting to the topic at hand is the association of entropy metrics to sleep depth (45,46).

## Hypotheses

Based on the above detailed theoretical background, the following hypotheses were formulated:

1. Recovery sleep following deprivation will be reflected by steeper spectral slope and higher spectral intercept compared to baseline levels (pre-sleep deprivation levels at corresponding times of day).
2. Slow wave activity will reflect the sleep pressure caused by sleep deprivation by increased values.
3. Normalized spectral entropy will report decreased complexity during recovery sleep.

## Methods Sample, Experimental Conditions & ECoG Recording

The sample consists of 9 mice (*Mus musculus, C57BL/6 and 129S4/SvJae hybrids*; 46), exposed to a 9 day-long continuous electrocorticograph (EcoG) recording containing a sleep deprivation paradigm with baseline, sleep deprivation, and recovery phases. At the time of surgery, the mice were 12 weeks old. Consecutive to implanting the electrodes, 5 days of habituation, then 9 days (Figure 1.) of recording was conducted. Each day consisted of an alternating light and dark cycle (with freely available food and water), namely, from 00:00 to 06:00 the lights were on, from 06:00 to 18:00 the lights were off and then on again from 18:00 to 24:00. The first day, serving as a baseline (BL), the mice were free roaming. Afterwards, for five days they were sleep deprived (sleep restriction days, SR) using a motorized wheel (14 cm in diameter, 5.8 cm in width, Lafayette Instrument, IN, USA) starting at 00:00 until 18:00, after which the mice were free roaming between 18:00 and 24:00. The last three days served as the recovery phase (recovery days, R), in which, following the same light and dark cycle, the mice were free roaming in their cages.

**Figure 1.**
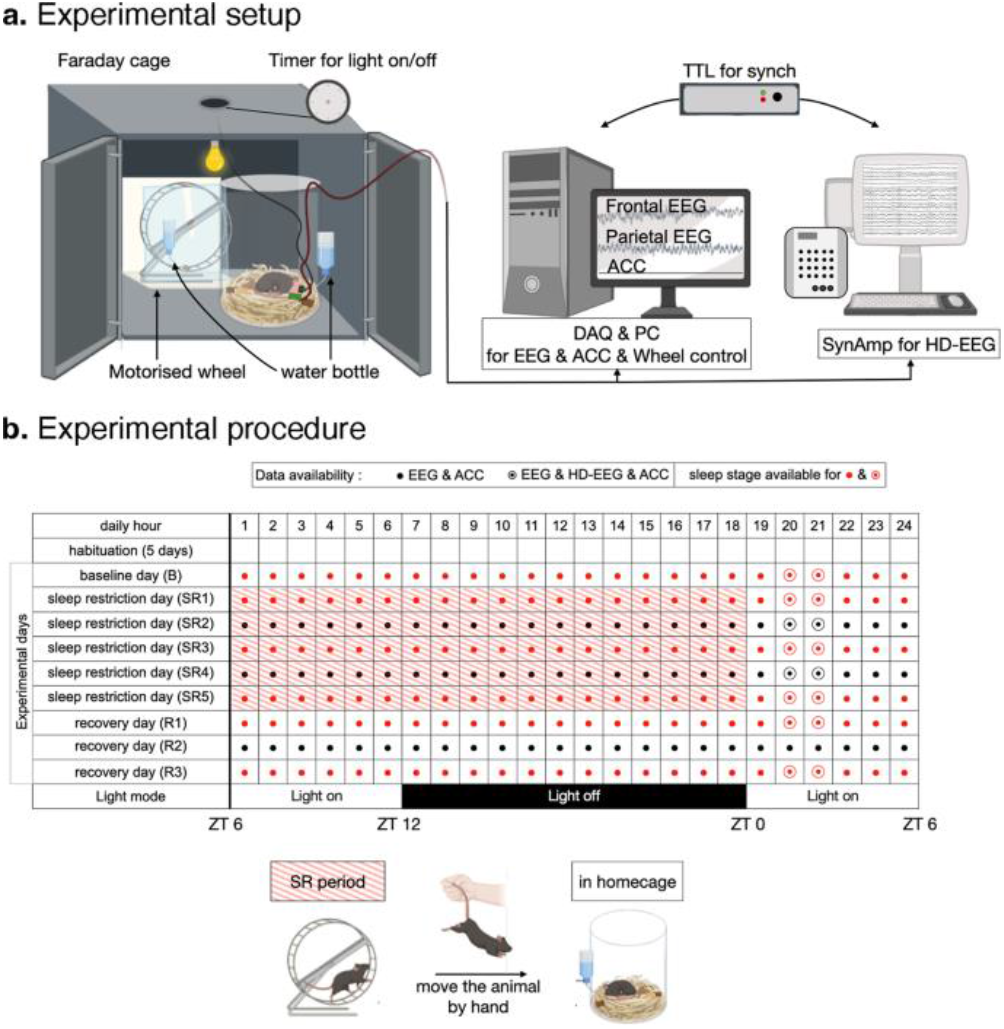
**Depiction of the experimental design**

Recordings were conducted in a sound-proof Faraday cage. The frontal and parietal registers were measured in a bipolar scheme referred to the left interparietal bone with ground electrode on the right interparietal bone. The signals were digitized after amplification (analog amplifier: QP511, Grass Technologies, West Warwick, RI, USA) using an analog-digital converter (Digidata 1440A, Molecular Devices, Sunnyvale, CA, USA). Sampling frequency was specified at 500 Hz, with a high-pass filter at 0.3 Hz, low-pass filter at 100 Hz, and 60 Hz hardware notch filter applied.

## Power spectrum

Power spectral density (PSD) was estimated using Welch’s method with a Hann window (n = 2000 point symmetric window, 4 sec), 50% overlap, and the number of discrete Fourier transform points used in the PSD estimate specified at 2000.

## Scale-free parameters of the power spectrum

The concept of scale-invariance (scale-free) is formalized such that

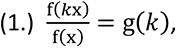

where *k* is a scaling factor, since the scaling does not depend on x. In practice this translates to “scaling the argument of the function is equivalent to a proportional scaling of the function itself” (7). This holds for the function f(x) = Cx^−α^, where α is the scaling exponent, essentially describing a power law, which is a density function in the form of p(x) = Cx^−α^.

Note that

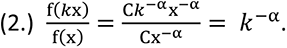

In the log-log space, an exponent is rendered a coefficient, transforming the function being investigated into a first-degree polynomial. A method, called fitting oscillations & one over f (FOOOF), used to extract the power-law exponent has been developed by Donoghue and colleges (17). It is a data-driven method with the purpose of describing the power spectrum on the assumption of the brain expressing scale-invariant, mono-fractal properties. It is mathematically expressed as:

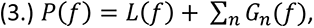

where P(f) is the sum of possibly multiple Gaussians Gn(f) and the aperiodic component L(f), which is parametrized as a Lorentzian function:

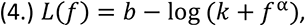

where b is the spectral intercept, α is the spectral slope, and k is the spectral knee. FOOOF computes the following parameters: 1) the scaling exponent (spectral slope); 2) the whitened peak, 3) width and 4) the expected value of multiple spectral peaks; 5) spectral intercept 6) and the spectral knee. The spectral exponent (also called spectral slope) refers to the background neural activity, the spectral peaks are the oscillatory activities superimposed over the background, the spectral intercept is the offset of the fitted curve, and the spectral knee is the point at which the power spectrum takes on its characteristic-coloured noise form on the logarithmically scaled space. The MATLAB implementation of the FOOOF fitting algorithm was applied over the frequency range of 0.5 to 48 Hz, with the following input parameters: width of peaks = default; maximum number of peaks = 0; minimum height of peaks = default; peak threshold = default; aperiodic mode = ‘knee’. The present study focuses on the spectral slope, the spectral intercept, and the spectral knee.

## Informational entropy-based characterization of neurophysiological signals

Entropy denotes the regularity of a signal – a highly ordered and therefore predictable signal is said to have low entropy, while an unpredictable and complex signal is characterized by high values of entropy. A measure of entropy, called Irregularity Index (II) has been proposed to measure EEG complexity (48). In practice, it involves calculating Shannon entropy (49) with log base 2 of the relative power of the time series signal, such that:

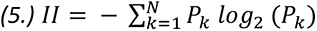

where k corresponds to the frequency λ_k_ and P_k_ is given by:

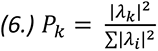

|λ_k_|^2^ being the power spectrum of a time-series at frequency λ_k_. However, this method lacks comparability – reflecting on this issue, namely that Shannon entropy does not yield a standardized measure, Zaccarelli and colleges (50) proposed and validated an index called Normalized Spectral Entropy (H_Sn_), normalized to the range of values between 0 and 1, such that:

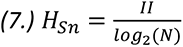

where II is the Irregularity Index (or spectral entropy) and N is the number of frequency bins. The normalisation ensures that the entropy value lies between 0 (representing a perfectly predictable or periodic signal) and 1 (representing a maximally random system with a flat spectrum). Important to note that in the framework of criticality, a maximally random system lacks structure, therefore should bear no information content and it should be assigned an entropy value of zero. We attempt to resolve this contradiction between this and our choice of Hsn by our motive to uniformalize our analysis and thus keep it in the frequency domain, and by previous empirical results; namely, that Spectral entropy has been used in drowsiness detection (51), it has been associated with sleep deprivation in humans (52). The present study utilizes the EntropyHub MATLAB toolbox (53).

## Artifact rejection & Data analysis

Artifact rejection was done manually, focusing on the harmonics of the network noise (60 Hz); in average, data loss was below 5%. The 24-hour long recording sessions were partitioned into one-hour long segments. Out of the 9 mice, subject 6 had faulty recordings on the frontal channel, therefore, sleep scoring is missing for this animal. Accordingly, subject 6 was omitted from the analysis. Sleep scoring (WAKE, REM, NREM) was done on day 1 (baseline ̶ BL), day 2 (sleep restriction ̶ SR1), day 4 (sleep restriction ̶ SR3), day 6 (sleep restriction ̶ SR5), day 7 (recovery ̶ R1), and day 9 (recovery ̶ R3). Due to missing data and inappropriately low time spent in NREM, the 18:00 to 19:00 time ranges of all subjects are omitted from the analyses, therefore our analysis focused on the 19:00 to 24:00 periods of the above specified days with associated sleep scoring information, considering NREM, REM, and WAKE episodes separately; in addition, a fourth analysis was performed on all non-artefactual segments, on an hourly basis, without considering sleep-wake staging information. Moreover, the parietal (Results section) and frontal (Appendix section) channels were independently analyzed. The data was analyzed using generalized linear models with two hierarchically organized repeated measure factors (DAYS: baseline, sleep restriction 1, sleep restriction 3, sleep restriction 5, sleep recovery 1, sleep recovery 3; HOURS: 19-20, 20-21, 21-22, 22- 23, 23-24). These hours were selected for targeting the immediate post-deprivation periods during the SR days, as well as the corresponding times-of-days on baseline and sleep recovery days. We chose this approach in order to maximize the sleep deprivation effects in our analyses. Significant main effects were further investigated by Bonferroni adjusted post-hoc tests. We considered α < 0.01 to be statistically significant. The analyzed metrics are the summed-up, natural logarithm transformed energy within the SWA frequency range of the power spectrum (0.75-4.5 Hz), the spectral slope, spectral intercept, spectral knee, and normalized spectral entropy.

## Results

### Time Spent in WAKEFULNESS, NREM and REM

Table 1.A, 1.B and 1.C shows the descriptive statistics (sample size, arithmetic mean, minimum, maximum, and standard deviation) of time spent in WAKE, NREM and REM phases, respectively, during the sleep scored days in each hour from 19:00 to 24:00. Effective hypothesis decomposition revealed that NREM episodes varied significantly over the experimental days (DAYS: F(5, 25) = 6.0827; p = .0004; η_p_^2^ = 0.46). More precisely, comparing BL to SR5 (t = -4.2907; p = .002), SR1 to SR5 (t = -4.1263; p = .003), and SR5 to R3 (t = 4.0011; p = .004) yielded significant results. In addition, statistically significant differences were observed in terms of time spent in wakefulness (WAKE) throughout the experimental days (Fig 2.; DAYS: F(5, 35) = 9.8923; p < .00001; η_p_ ^2^ = 0.59). Bonferroni corrected post-hoc tests of the DAYS independent variable revealed statistically significant differences contrasting BL sleep to SR3 (t = 3.561; p = .0163) and SR5 (t = 6.0363; p < .001), SR1 to SR5 (t = 4.7572; p < .001), and SR5 to R3 (t = -5.2884; p < .001). Considering REM phase, similarly, only the DAYS independent variable was significant (DAYS: F(5, 35) = 7.6155; p < .001; η_p_ ^2^ = 0.52). Post-hoc test showed significant differences on the following comparison: BL vs. SR1 (t = -3.7864; p = .0086), SR3 (t = -3.6622; p = .0123) and SR5 (t = -4.7367; p < .001); SR1 vs. R1 (t = 3.2528; p = 0.038); and SR5 vs. R1 (t = 4.2031; p = .0026) and R3 (t = 3.2455; p = .0387).

**Figure 2.**
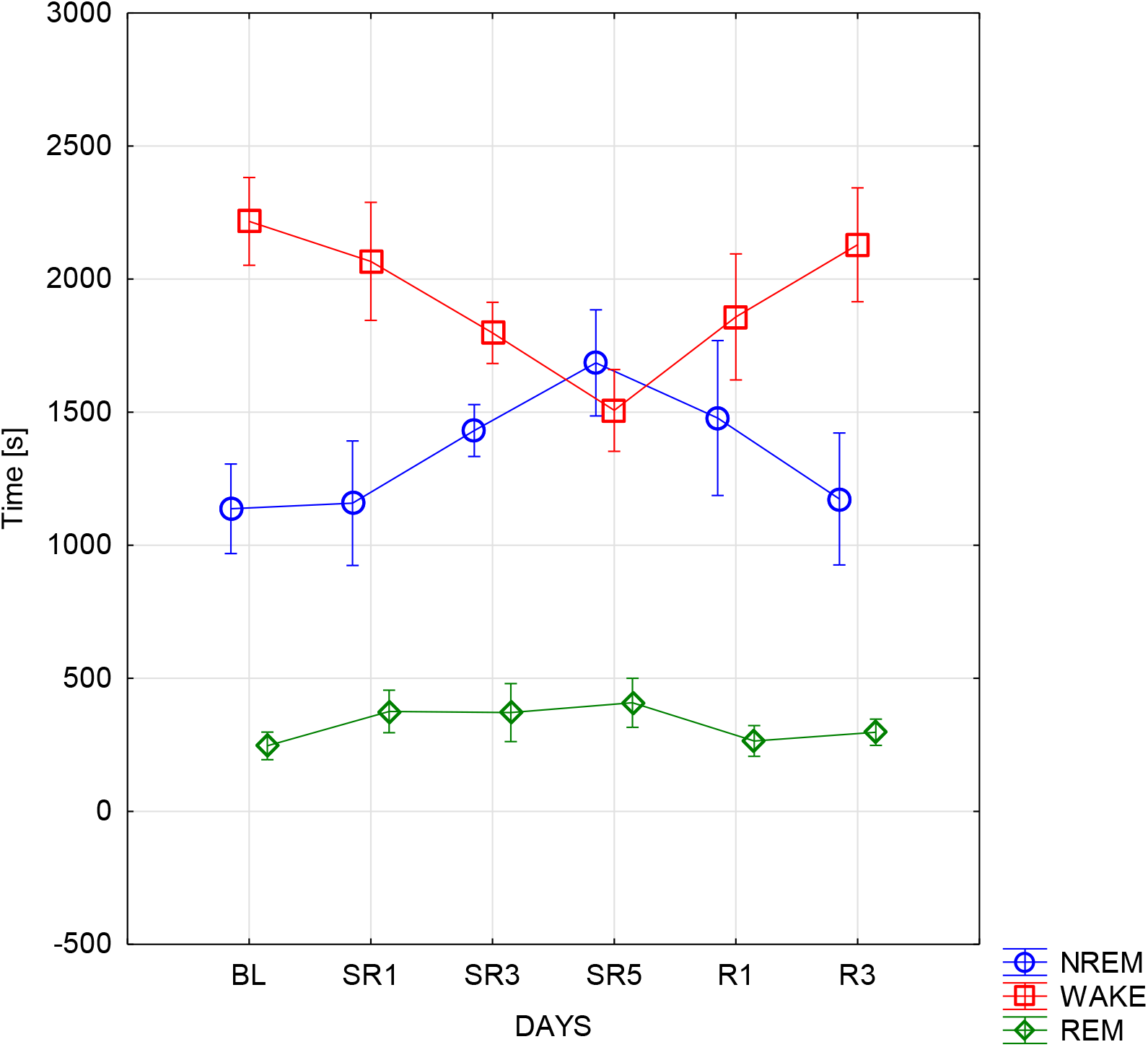
**Time spent in distinct behavioural states (WAKEFULNESS, NREM and REM) as functions of the experimental intervention. Note the exclusive focus on hours 19:00-24:00 (see details in the section Methods).** note: Error bars denote standard errors; BL = Baseline, SR = Sleep Restriction, R = Recovery

**Table 1.** A Descriptive statistics of time (min) spent in NREM.

|  | 18:00-19:00 | 19:00-20:00 | 20:00-21:00 | 21:00-22:00 | 22:00-23:00 | 23:00-24:00 |
| --- | --- | --- | --- | --- | --- | --- |
| **Baseline** | 25.83(11.42) | 18.46(6.74) | 19.83(7.16) | 20.19(6.38) | 17.17(8.87) | 19.1(8.79) |
| **Sleep Restriction 1** | 35.73(14.83) | 16.79(9.26) | 21.83(6.91) | 22.4(13.51) | 13.71(9.97) | 21.77(10.4) |
| **Sleep Restriction 3** | 45.69(12.49) | 27.81(10.73) | 28.06(9.62) | 20.54(8.85) | 16.33(7.08) | 26.48(8.44) |
| **Sleep Restriction 5** | 52.96(7.75) | 33.98(11.96) | 30.98(12.96) | 27.31(8.22) | 26.83(13.82) | 21.33(10.98) |
| **Sleep Recovery 1** | 32.27(16.16) | 21.79(9.88) | 25.04(13.01) | 29.4(11.62) | 20.58(11.45) | 26.35(11.69) |
| **Sleep Recovery 3** | 27.33(15.64) | 18.02(8.97) | 20.88(12.9) | 23.02(10.14) | 16.04(3.62) | 19.88(8.07) |

**Table 1.** B Descriptive statistics of time (min) spent in WAKE.

|  | 18:00-19:00 | 19:00-20:00 | 20:00-21:00 | 21:00-22:00 | 22:00-23:00 | 23:00-24:00 |
| --- | --- | --- | --- | --- | --- | --- |
| **Baseline** | 31.04 (9.99) | 38.33 (6.53) | 35.72 (7.23) | 35.67 (5.52) | 38.25 (7.97) | 36.77 (7.32) |
| **Sleep Restriction 1** | 19.12 (12.22) | 36.58 (8.35) | 33.81 (6.08) | 31.9 (11.71) | 37.77 (7.62) | 32.12 (9.13) |
| **Sleep Restriction 3** | 11.87 (11.48) | 26.06 (9.99) | 26.88 (9.33) | 33.16 (6.47) | 35.68 (4.02) | 28.04 (7.25) |
| **Sleep Restriction 5** | 8.13 (5.93) | 20.58 (9.49) | 22.75 (8.97) | 25.70 (5.42) | 25.75 (11.45) | 30.77 (8.49) |
| **Sleep Recovery 1** | 25.042 (14.53) | 34.06 (9.07) | 30.75 (11.2) | 26.77 (9.37) | 34.17 (10.37) | 29.06 (9.93) |
| **Sleep Recovery 3** | 29.14 (14.14) | 37.02 (7.89) | 34.38 (10.5) | 32.39 (8.52) | 38.75 (3.54) | 34.88 (7.04) |

**Table 1.**
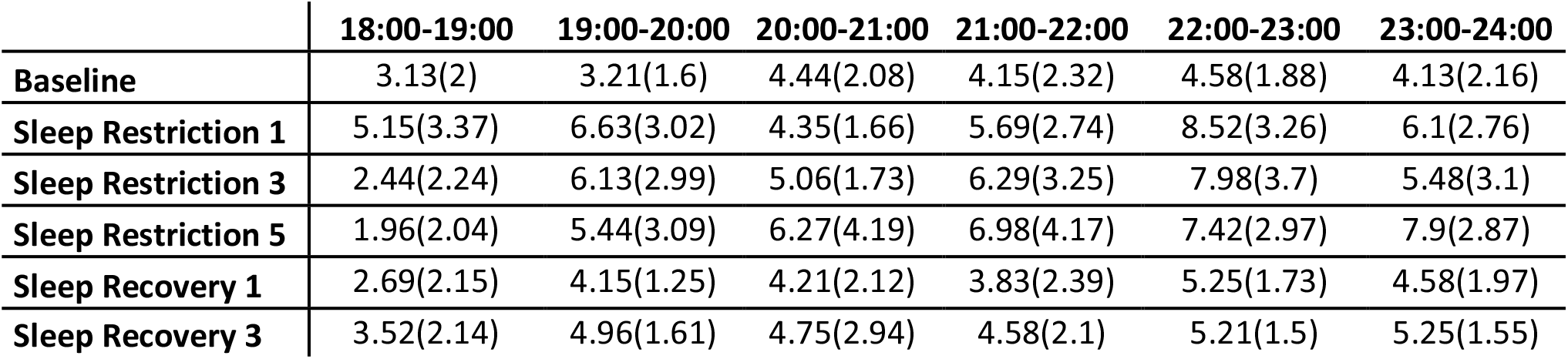
C Descriptive statistics of time (min) spent in REM.

In other words, the mice appeared to spend more time in NREM and REM sleep as a response to sleep deprivation, which decrement returned to baseline values during recovery days. On the other hand, WAKE appeared to follow an inverse pattern, that is, it increased during sleep deprived days, then renormalized.

### Slow Wave Activity (SWA)

In terms of SWA no significant effects of sleep deprivation or recovery hour emerged for WAKE (Table 2.A) or REM (Table 2.C), whereas for NREM (Table 2.B) scored segments, both independent variables (DAYS: F(5, 35) = 3.6227; p = .00957; η_p_ ^2^ = 0.34, HOURS: F(4, 28) = 10.04; p = .00004; η_p_ ^2^ = 0.58) and their interaction (F(20, 140) = 2.0156; p = .00983; η_p_^2^ = 0.22) were significant (Figure 3).

**Figure 3.**
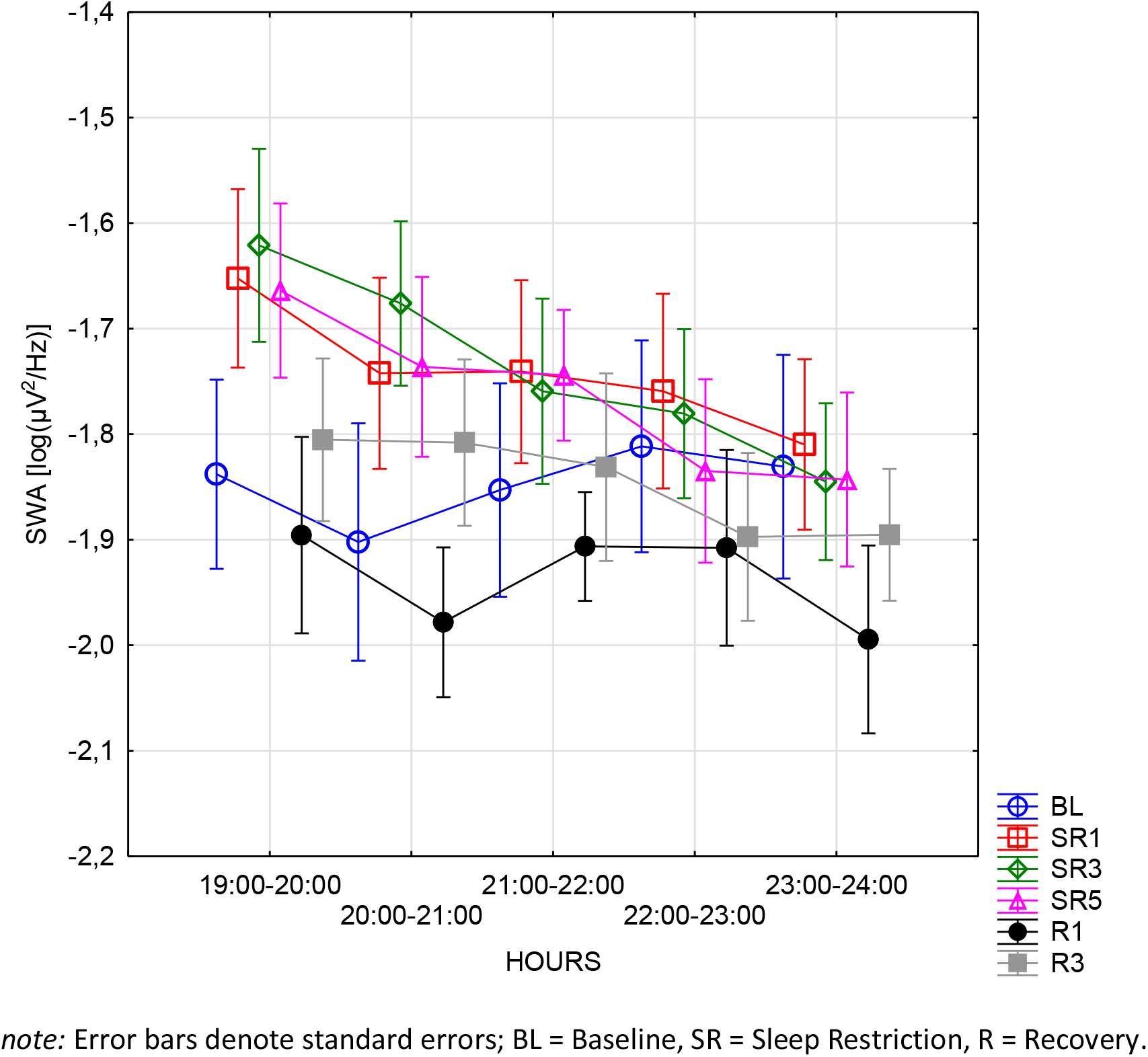
**NREM sleep ECoG SWA as a function of the experimental intervention and time of day**

**Table 2.**
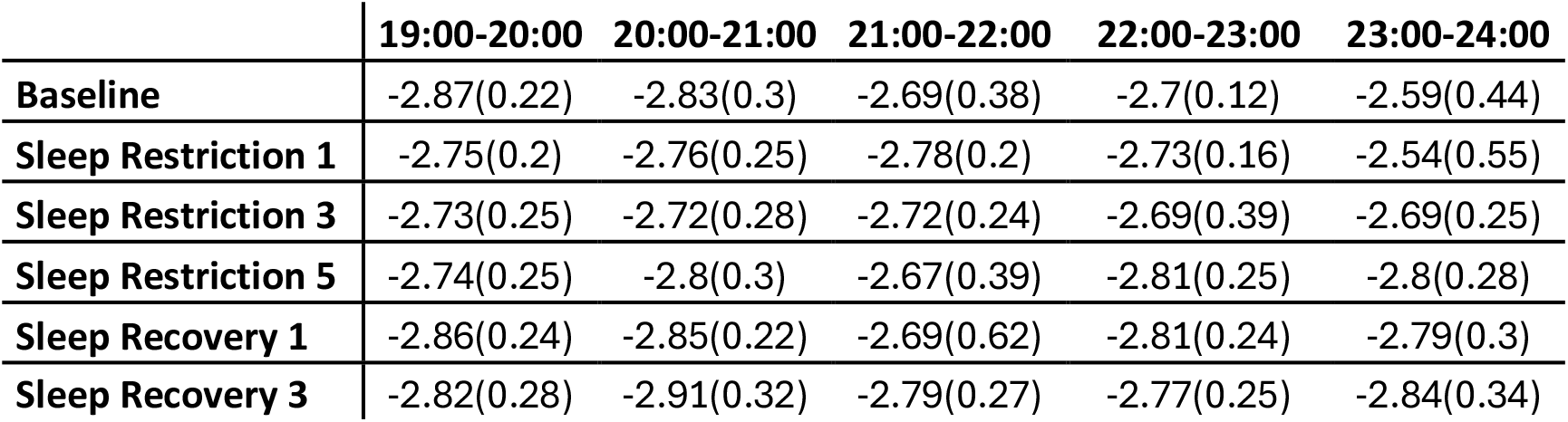
A Descriptive statistics of SWA in WAKE.

**Table 2.** B Descriptive statistics of SWA in NREM.

|  | 19:00-20:00 | 20:00-21:00 | 21:00-22:00 | 22:00-23:00 | 23:00-24:00 |
| --- | --- | --- | --- | --- | --- |
| **Baseline** | -1.84(0.25) | -1.9(0.32) | -1.85(0.29) | -1.81(0.28) | -1.83(0.3) |
| **Sleep Restriction 1** | -1.65(0.24) | -1.74(0.26) | -1.74(0.25) | -1.76(0.26) | -1.81(0.23) |
| **Sleep Restriction 3** | -1.62(0.26) | -1.68(0.22) | -1.76(0.25) | -1.78(0.23) | -1.84(0.21) |
| **Sleep Restriction 5** | -1.66(0.23) | -1.74(0.24) | -1.74(0.18) | -1.83(0.25) | -1.84(0.23) |
| **Sleep Recovery 1** | -1.9(0.26) | -1.98(0.2) | -1.91(0.15) | -1.91(0.26) | -1.99(0.25) |
| **Sleep Recovery 3** | -1.81(0.22) | -1.81(0.22) | -1.83(0.25) | -1.9(0.22) | -1.9(0.18) |

**Table 2.** C Descriptive statistics of SWA in REM.

|  | 19:00-20:00 | 20:00-21:00 | 21:00-22:00 | 22:00-23:00 | 23:00-24:00 |
| --- | --- | --- | --- | --- | --- |
| **Baseline** | -2.74(0.26) | -2.79(0.34) | -2.8(0.24) | -2.71(0.2) | -2.7(0.24) |
| **Sleep Restriction 1** | -2.64(0.3) | -2.73(0.27) | -2.56(0.32) | -2.74(0.22) | -2.71(0.27) |
| **Sleep Restriction 3** | -2.69(0.28) | -2.73(0.29) | -2.86(0.27) | -2.77(0.25) | -2.94(0.21) |
| **Sleep Restriction 5** | -2.73(0.24) | -2.75(0.26) | -2.74(0.16) | -2.77(0.2) | -2.78(0.23) |
| **Sleep Recovery 1** | -2.75(0.27) | -2.88(0.14) | -2.82(0.2) | -2.93(0.19) | -2.84(0.13) |
| **Sleep Recovery 3** | -2.75(0.24) | -2.77(0.16) | -2.77(0.17) | -2.82(0.21) | -2.73(0.27) |

**Table 2.** D Descriptive statistics of SWA in ALL recorded periods.

|  | 19:00-20:00 | 20:00-21:00 | 21:00-22:00 | 22:00-23:00 | 23:00-24:00 |
| --- | --- | --- | --- | --- | --- |
| **Baseline** | -2.09(0.24) | -2.18(0.29) | -2.12(0.24) | -2.06(0.18) | -2.06(0.29) |
| **Sleep Restriction 1** | -1.95(0.23) | -2.06(0.19) | -2.11(0.28) | -2.03(0.19) | -2.06(0.26) |
| **Sleep Restriction 3** | -2.06(0.18) | -2.12(0.23) | -2.08(0.26) | -2.06(0.21) | -2.22(0.16) |
| **Sleep Restriction 5** | -2.24(0.22) | -2.25(0.36) | -2.18(0.17) | -2.24(0.24) | -2.2(0.25) |
| **Sleep Recovery 1** | -2.2(0.27) | -2.33(0.21) | -2.28(0.33) | -2.23(0.2) | -2.35(0.31) |
| **Sleep Recovery 3** | -2.07(0.18) | -2.17(0.29) | -2.17(0.21) | -2.13(0.18) | -2.19(0.23) |

*note:* Table 2.A, 2.B, 2.C and 2.D contains the descriptive statistics of SWA in WAKE, NREM, REM episodes, and when all scores were considered (ALL), in the following format: MEAN (STANDARD DEVIATION).

Throughout the experimental days, the following comparisons were significant: SR1 vs. R1 (t = 3.3424; p = .0298; d = 0.8082) and SR3 vs. R1 (t = 3.4187; p = .0241; d = 0.8267). Post-hoc tests of the HOURS independent variable showed significant differences contrasting the 19-20 vs 22-23 (t = 4.263; p = .0021; d = 0.3547) and 23-24 (t = 6.1433; p < .001; d = 0.5111) intervals, and the 20-21 vs 23-24 (t = 3.1069; p = .043; d = 0.2585) and 21-22 vs 23-24 (t = 3.1755; p = .0362; d = 0.2642) intervals. Post-hoc tests of the interaction term are reported in the Appendix. However, when all the scores (NREM, REM, WAKE; Table 2.D) were considered, only the DAYS independent variable was found to be statistically significant (F(5, 35) = 5.1116; p = .00127; η_p_ ^2^ = 0.42). Post-hoc tests revealed significant differences between BL and R1 (t = 3.2623; p = .037; d = 0.7235), and SR1 vs. R1 (t = 3.3408; p = .0299; d = 0.7409) and R3 (t = 4.3742; p = .0016; d = 0.9701). That is to say, ECoG power in the low frequency range is enhanced as a result of sleep deprivation, but sleep deprivation-related effects are mainly present during the NREM phase of the first few hours of sleep, which progressively become normalized (Figure 3).

### Spectral Intercept

In terms of the spectral intercept during WAKE (Table 3.A) periods only the DAYS independent variables was significant (F(5, 35) = 3.772; p = .0078; η_p_ ^2^ = 0.35); which according to Bonferroni corrected post-hoc test, indicated the difference between SR1 and R1 (t = 3.9772; p = .0051; d = 0.3632). However, for REM (Table 3.C, Figure 4.B) multiple significant main effects were found (DAYS: F(5, 20) = 6.3814; p = .0011; η_p_ ^2^ = 0.61, HOURS: F(4, 16) = 18.7921; p < .001; η_p_ ^2^ = 0.82). Post-hoc tests showed significant comparisons for SR1 vs. R1 (t = 4.5644; p = .0028, d = 0.6934), SR3 vs. R1 (t = 3.5562; p = .0297; d = 0.5402) and SR5 vs. R1 (t = 4.4309; p = .0039; d = 0.406); and for 19-20 vs. 21-22 (t = 5.494; p < .001; d = 0.2924), 22-23 (t = 6.3141; p < .001; d = 0.3361) and 23-24 (t = 7.6253; p < .001; d = 0.4058), and 20-21 vs. 22-23 (t = 3.6356; p = .0223; d = 0.1935) and 23-24 (t = 4.9468; p = .0015; d = 0.2633). In addition, for NREM (Table 3.B, Figure 4.A), both independent variables (DAYS: F(5, 35) = 8.6165; p < .001; η_p_ ^2^ = 0.55, HOURS: F(4, 28) = 5.8956; p = .0014; η_p_ ^2^ = 0.45) and their interaction were found to be significant (Figure 4 ;F(20, 140) = 2.7777; p < .001; η_p_ ^2^ = 0.28). Post-hoc tests revealed significant differences between SR1 and R1 (t = 5.1768; p < .001; d = 1.0557) and R3 (t = 3.697; p = .0111; d = 0.7539), between SR3 and R1 (t = 4.3998; p = .0015; d = 0.8973), and finally between SR5 and R1 (t = 4.8638; p < .001; 0.9919). Lastly, for the aggregated segments (Table 3.D; WAKE, NREM, and REM included), significant differences were only observed throughout the experimental days (DAYS: F(5, 35) = 8.8427; p < .001; η_p_ ^2^ = 0.56). More precisely, BL vs. R1 (t = 4.5301; p < .001; d = 0.6801), SR1 vs. SR5 (t = 3.5787; p = .0153; d = 0.5372) and R1 (t = 6.3422; p < .001; d = 0.9521), SR3 vs. R1 (t = 3.8676; p = .0068; d = 0.5805).

**Figure 4.**
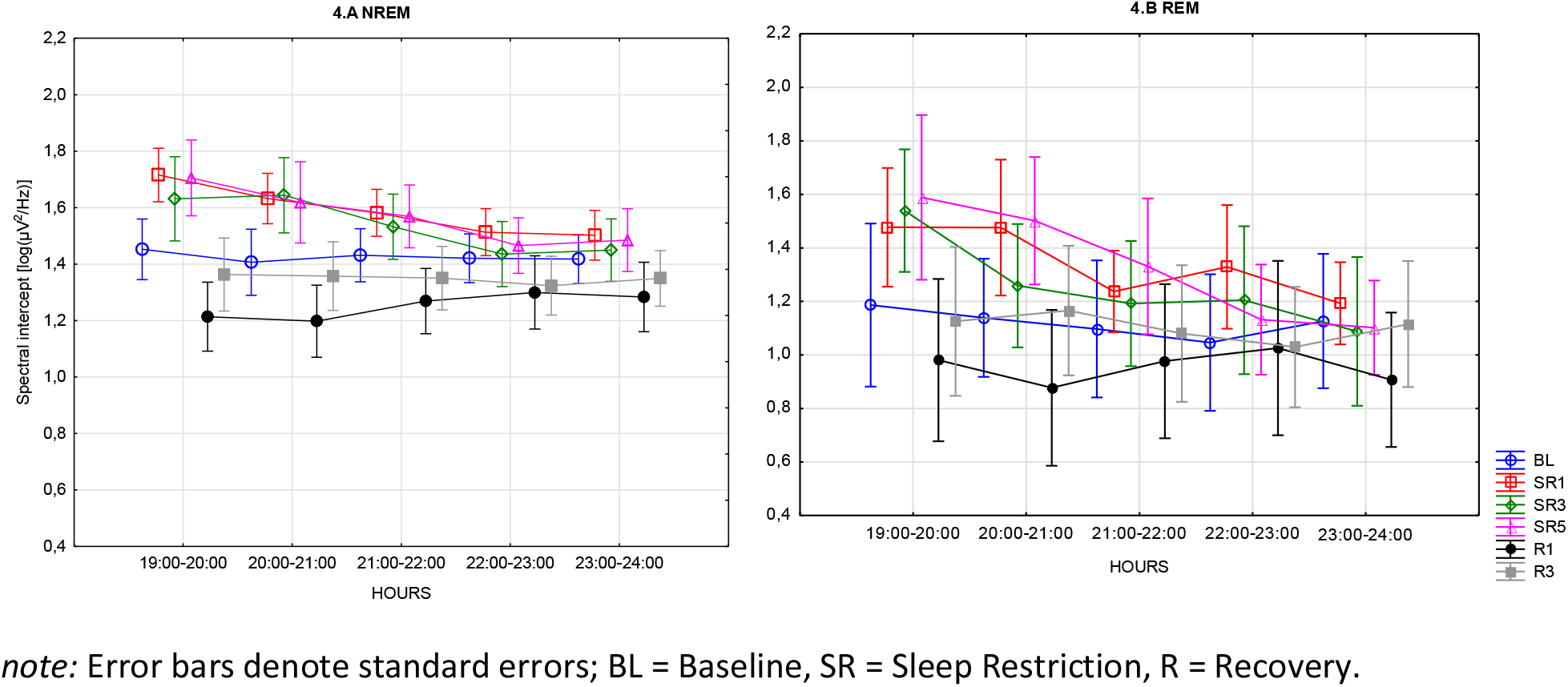
**Sleep ECoG spectral intercepts as functions of the experimental intervention and time of day. A. NREM sleep ECoG spectral intercepts. B. REM sleep ECoG spectral intercepts**

**Table 3.** A Descriptive statistics of Spectral Intercept in WAKE.

|  | 19:00-20:00 | 20:00-21:00 | 21:00-22:00 | 22:00-23:00 | 23:00-24:00 |
| --- | --- | --- | --- | --- | --- |
| **Baseline** | -0.32(0.8) | -0.27(0.71) | -0.18(0.78) | -0.18(0.65) | -0.24(0.66) |
| **Sleep Restriction 1** | -0.02(0.61) | -0.12(0.67) | -0.15(0.68) | 0.14(0.6) | -0.23(0.45) |
| **Sleep Restriction 3** | -0.17(0.65) | -0.16(0.71) | -0.13(0.68) | -0.09(0.7) | -0.29(0.58) |
| **Sleep Restriction 5** | -0.21(0.73) | -0.18(0.65) | -0.15(0.69) | -0.18(0.64) | 0.01(0.64) |
| **Sleep Recovery 1** | -0.38(0.74) | -0.35(0.76) | -0.42(0.61) | -0.2(0.64) | -0.25(0.71) |
| **Sleep Recovery 3** | -0.14(0.69) | -0.33(0.66) | -0.2(0.56) | -0.22(0.69) | -0.2(0.71) |

**Table 3.** B Descriptive statistics of Spectral Intercept in NREM.

|  | 19:00-20:00 | 20:00-21:00 | 21:00-22:00 | 22:00-23:00 | 23:00-24:00 |
| --- | --- | --- | --- | --- | --- |
| **Baseline** | 1.45(0.3) | 1.41(0.33) | 1.43(0.27) | 1.42(0.24) | 1.42(0.24) |
| **Sleep Restriction 1** | 1.72(0.27) | 1.63(0.25) | 1.58(0.24) | 1.51(0.24) | 1.5(0.25) |
| **Sleep Restriction 3** | 1.63(0.42) | 1.64(0.38) | 1.53(0.33) | 1.44(0.33) | 1.45(0.31) |
| **Sleep Restriction 5** | 1.71(0.38) | 1.62(0.41) | 1.57(0.31) | 1.47(0.28) | 1.49(0.32) |
| **Sleep Recovery 1** | 1.21(0.35) | 1.2(0.36) | 1.27(0.33) | 1.3(0.37) | 1.28(0.35) |
| **Sleep Recovery 3** | 1.36(0.37) | 1.36(0.34) | 1.35(0.32) | 1.32(0.29) | 1.35(0.28) |

**Table 3.**
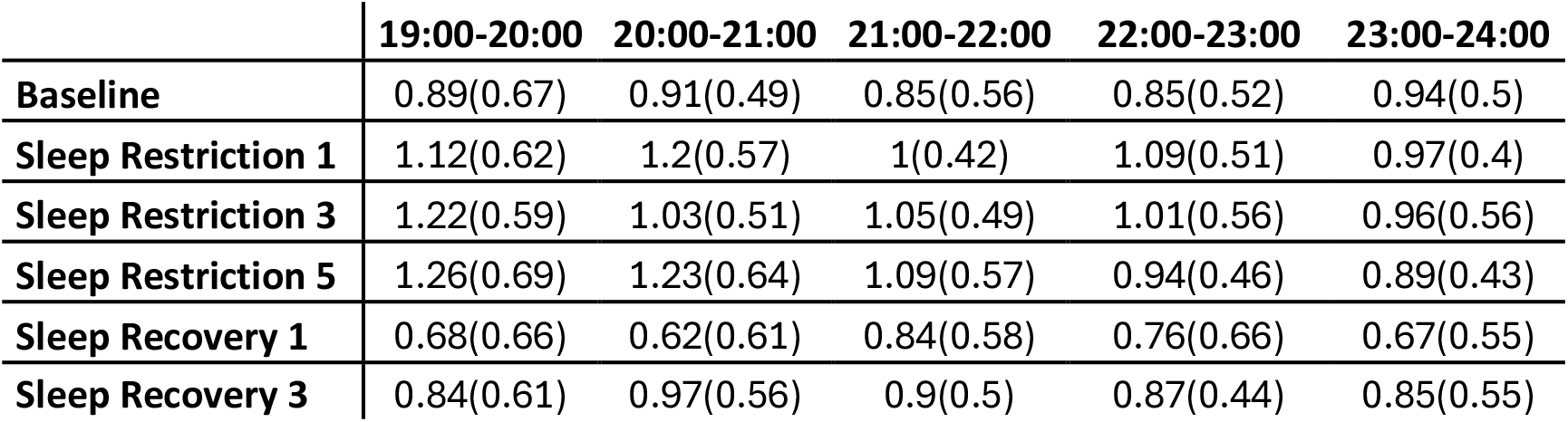
C Descriptive statistics of Spectral Intercept in REM.

|  | 19:00-20:00 | 20:00-21:00 | 21:00-22:00 | 22:00-23:00 | 23:00-24:00 |
| --- | --- | --- | --- | --- | --- |
| **Baseline** | 0.89(0.67) | 0.91(0.49) | 0.85(0.56) | 0.85(0.52) | 0.94(0.5) |
| **Sleep Restriction 1** | 1.12(0.62) | 1.2(0.57) | 1(0.42) | 1.09(0.51) | 0.97(0.4) |
| **Sleep Restriction 3** | 1.22(0.59) | 1.03(0.51) | 1.05(0.49) | 1.01(0.56) | 0.96(0.56) |
| **Sleep Restriction 5** | 1.26(0.69) | 1.23(0.64) | 1.09(0.57) | 0.94(0.46) | 0.89(0.43) |
| **Sleep Recovery 1** | 0.68(0.66) | 0.62(0.61) | 0.84(0.58) | 0.76(0.66) | 0.67(0.55) |
| **Sleep Recovery 3** | 0.84(0.61) | 0.97(0.56) | 0.9(0.5) | 0.87(0.44) | 0.85(0.55) |

**Table 3.** D Descriptive statistics of Spectral Intercept in ALL recorded periods.

|  | 19:00-20:00 | 20:00-21:00 | 21:00-22:00 | 22:00-23:00 | 23:00-24:00 |
| --- | --- | --- | --- | --- | --- |
| **Baseline** | 0.9(0.52) | 0.83(0.42) | 0.88(0.44) | 0.96(0.26) | 0.86(0.29) |
| **Sleep Restriction 1** | 1.22(0.33) | 1.01(0.38) | 0.94(0.45) | 1.12(0.34) | 0.82(0.35) |
| **Sleep Restriction 3** | 0.84(0.57) | 0.82(0.41) | 0.93(0.54) | 0.95(0.51) | 0.66(0.47) |
| **Sleep Restriction 5** | 0.64(0.77) | 0.71(0.75) | 0.77(0.54) | 0.79(0.44) | 0.88(0.42) |
| **Sleep Recovery 1** | 0.6(0.6) | 0.47(0.58) | 0.41(0.54) | 0.74(0.44) | 0.55(0.56) |
| **Sleep Recovery 3** | 0.87(0.56) | 0.72(0.61) | 0.72(0.37) | 0.87(0.42) | 0.81(0.41) |

note: Table 3.A, 2.B, 2.C and 2.D contains the descriptive statistics of Spectral Intercept in WAKE, NREM, REM episodes, and when all scores were considered (ALL), in the following format: MEAN (STANDARD DEVIATION).

### Spectral Slope

The spectral slopes of the ECoG records measured during the WAKE (Table 4.A) segments did not show significant sleep deprivation-related effects. On the other hand, during REM (Table 4.C, Figure 5.B) segments, both main effects (DAYS: F(5,20) = 6.4964; p = .00097; η_p_^2^ = 0.61, HOURS: F(4, 16) = 15.167; p = .00003; η_p_^2^ = 0.79) were significant, however, their interaction (F(20, 80) = 2.0016; p = .01576; η_p_^2^ = 0.33) only marginally. More precisely, the comparisons of SR1 vs. R1 (t = 4.5026; p = .0033; d = 0.7169), SR3 vs. R1 (t = 3.5442; p = .0305; d = 0.5643), and SR5 vs R1 (t = 4.4349; p = .0038; d = 0.7061) were found to be significant. Post-hoc tests of the HOURS independent variable also revealed significant differences: 19-20 vs. 21-22 (t = 5.0856; p = .0011; d = 0.2904), 22-23 (t = 5.5744; p > .001; d = 0.3183), 23-24 (t = 6.6224; p < .001; d = 0.3782), and 20-21 vs. 23-24 (t = 4.5959; p = .0029; d = 0.2625). An analysis with identical parameters was repeated for the NREM (Table 4.B, Figure 5.A) segments, which revealed significant main effects of DAYS (F(5, 35) = 10.86; p < .00001; η_p_^2^ = 0.61), HOURS (F(4, 28) = 4.3159; p = .00762; η_p_^2^ = 0.38) and their interaction (F(20, 140) = 2.5704; p = .00069; η_p_^2^ = 0.27). Post-hoc tests revealed significant differences between BL and R1 (t = 3.3979; p = .00256; d = 0.6311), SR1 and R1 (t = 6.0085; p < .001; d = 1.1159), R3 (t = 4.1518; p = .0031; d = 0.7711), SR3 and R1 (t = 5.2516; p < .001; d = 0.9753), R3 (t = 3.3948; p = .0258; d = 0.6305), and lastly, between SR5 and R1 (t = 5.2073; p < .001; d = 0.9671), R3 (t = 3.3505; p = .0291;d = 0.6305).

**Figure 5.**
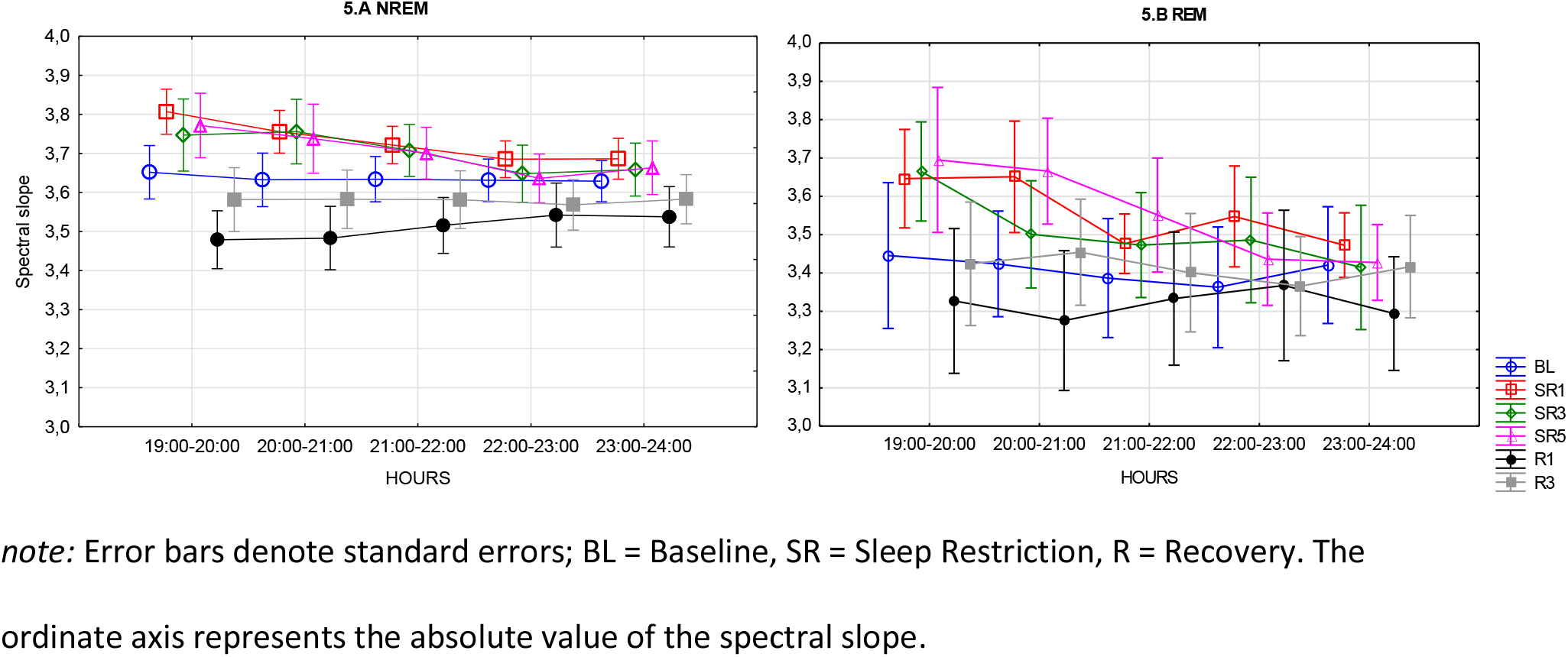
**Sleep ECoG spectral slopes as functions of the experimental intervention and times of day. A. NREM sleep ECoG spectral slopes. B. REM sleep ECoG spectral slopes**

**Table 4.** A Descriptive statistics of Spectral Slope in WAKE.

|  | 19:00-20:00 | 20:00-21:00 | 21:00-22:00 | 22:00-23:00 | 23:00-24:00 |
| --- | --- | --- | --- | --- | --- |
| **Baseline** | 2.43(0.55) | 2.47(0.47) | 2.53(0.54) | 2.54(0.44) | 2.48(0.43) |
| **Sleep Restriction 1** | 2.65(0.4) | 2.58(0.44) | 2.55(0.44) | 2.76(0.43) | 2.51(0.29) |
| **Sleep Restriction 3** | 2.54(0.41) | 2.55(0.45) | 2.57(0.42) | 2.61(0.46) | 2.47(0.36) |
| **Sleep Restriction 5** | 2.5(0.48) | 2.54(0.42) | 2.56(0.44) | 2.53(0.4) | 2.69(0.45) |
| **Sleep Recovery 1** | 2.4(0.49) | 2.41(0.5) | 2.33(0.38) | 2.5(0.41) | 2.48(0.46) |
| **Sleep Recovery 3** | 2.56(0.45) | 2.41(0.43) | 2.5(0.34) | 2.49(0.46) | 2.52(0.46) |

**Table 4.**
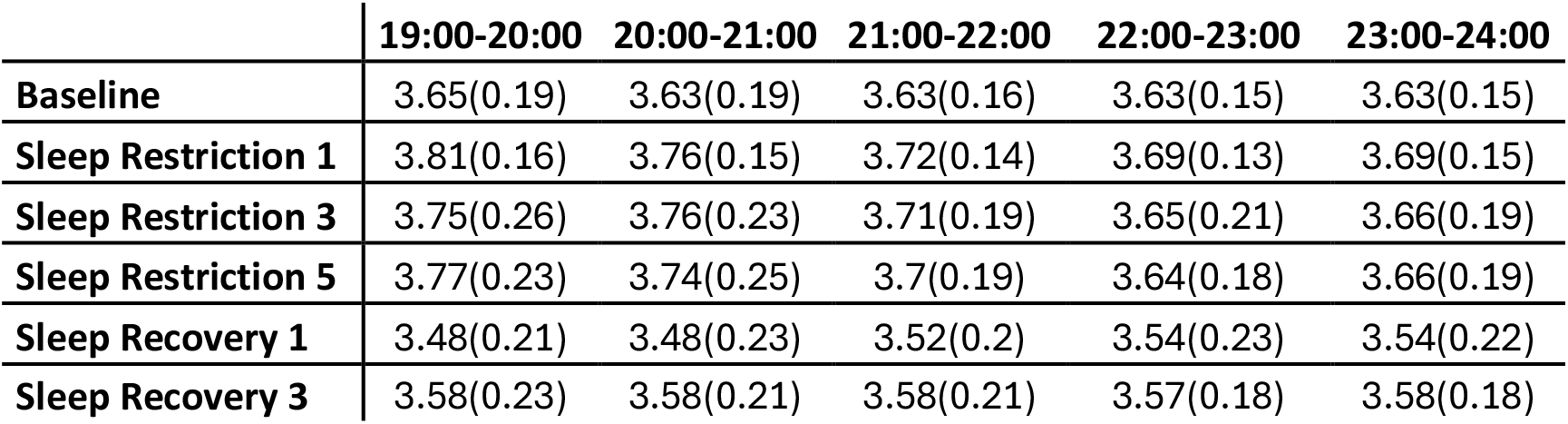
B Descriptive statistics of Spectral Slope in NREM.

|  | 19:00-20:00 | 20:00-21:00 | 21:00-22:00 | 22:00-23:00 | 23:00-24:00 |
| --- | --- | --- | --- | --- | --- |
| **Baseline** | 3.65(0.19) | 3.63(0.19) | 3.63(0.16) | 3.63(0.15) | 3.63(0.15) |
| **Sleep Restriction 1** | 3.81(0.16) | 3.76(0.15) | 3.72(0.14) | 3.69(0.13) | 3.69(0.15) |
| **Sleep Restriction 3** | 3.75(0.26) | 3.76(0.23) | 3.71(0.19) | 3.65(0.21) | 3.66(0.19) |
| **Sleep Restriction 5** | 3.77(0.23) | 3.74(0.25) | 3.7(0.19) | 3.64(0.18) | 3.66(0.19) |
| **Sleep Recovery 1** | 3.48(0.21) | 3.48(0.23) | 3.52(0.2) | 3.54(0.23) | 3.54(0.22) |
| **Sleep Recovery 3** | 3.58(0.23) | 3.58(0.21) | 3.58(0.21) | 3.57(0.18) | 3.58(0.18) |

**Table 4.** C Descriptive statistics of Spectral Slope in REM.

|  | 19:00-20:00 | 20:00-21:00 | 21:00-22:00 | 22:00-23:00 | 23:00-24:00 |
| --- | --- | --- | --- | --- | --- |
| **Baseline** | 3.26(0.42) | 3.28(0.31) | 3.22(0.36) | 3.23(0.33) | 3.3(0.31) |
| **Sleep Restriction 1** | 3.42(0.38) | 3.48(0.34) | 3.33(0.24) | 3.4(0.3) | 3.33(0.25) |
| **Sleep Restriction 3** | 3.46(0.35) | 3.36(0.31) | 3.37(0.3) | 3.35(0.35) | 3.33(0.33) |
| **Sleep Restriction 5** | 3.48(0.44) | 3.48(0.4) | 3.39(0.35) | 3.31(0.28) | 3.28(0.26) |
| **Sleep Recovery 1** | 3.13(0.42) | 3.1(0.39) | 3.23(0.36) | 3.19(0.41) | 3.13(0.34) |
| **Sleep Recovery 3** | 3.23(0.38) | 3.32(0.34) | 3.27(0.32) | 3.26(0.27) | 3.24(0.33) |

**Table 4.** D Descriptive statistics of Spectral Slope in ALL recorded periods.

|  | 19:00-20:00 | 20:00-21:00 | 21:00-22:00 | 22:00-23:00 | 23:00-24:00 |
| --- | --- | --- | --- | --- | --- |
| **Baseline** | 3.26(0.36) | 3.23(0.27) | 3.25(0.3) | 3.31(0.18) | 3.24(0.18) |
| **Sleep Restriction 1** | 3.47(0.22) | 3.33(0.25) | 3.28(0.29) | 3.42(0.23) | 3.22(0.22) |
| **Sleep Restriction 3** | 3.2(0.4) | 3.2(0.25) | 3.29(0.34) | 3.31(0.34) | 3.12(0.29) |
| **Sleep Restriction 5** | 3.06(0.52) | 3.13(0.49) | 3.16(0.36) | 3.17(0.29) | 3.26(0.29) |
| **Sleep Recovery 1** | 3.05(0.4) | 2.98(0.39) | 2.91(0.36) | 3.15(0.28) | 3.02(0.38) |
| **Sleep Recovery 3** | 3.24(0.39) | 3.14(0.4) | 3.13(0.23) | 3.24(0.28) | 3.21(0.27) |

note: Table 4.A, 2.B, 2.C and 2.D contains the descriptive statistics of Spectral Slope in WAKE, NREM, REM episodes, and when all scores were considered (ALL), in the following format: MEAN (STANDARD DEVIATION).

Furthermore, post-hoc testing of the HOURS independent variable revealed the 19-20 vs. 22-23 (t = 3.5815; p = .0127; d = 0.2772) comparison to be marginally significant. Last, but not least, the aggregated segments (Table 4.D; unifying WAKE, NREM and REM) did indicate significant differences only for the DAYs independent variable (F(5, 25) = 9.3007; p < .001; η_p_^2^ = 0.5706). Significant post-hoc comparisons are the following: BL vs. SR1 (t = 4.6894; p < .001; d = 0.7219), SR1 vs. SR5 (t = 3.7998; p = .0083; d = 0.5849) and R1 (t = 6.4275; p < .001; d = 0.9894), SR3 vs. R1 (t = 3.9846; p = .0049; d = 0.6134). In other words, the power spectrum appeared to respond with enhanced slow to high frequency ratios as captured by the steeper fitted curves to the ECoG spectra of sleep periods following sleep deprivation (SR1) relative to baseline sleep (BL).

### Spectral Knee

Table 5.A-C details the descriptives of the spectral knee, that is, the point at which the log-log power spectrum linearizes. It appears that the spectral knee is shifted to the right as opposed to human data (Figure 7.A and 7.B). More precisely, it averages around 9 Hz with a range of 7 to 11 Hz in mice. As a result of the substantial right shift, the spectral slope is unfit to describe slow oscillations.

**Figure 6.**
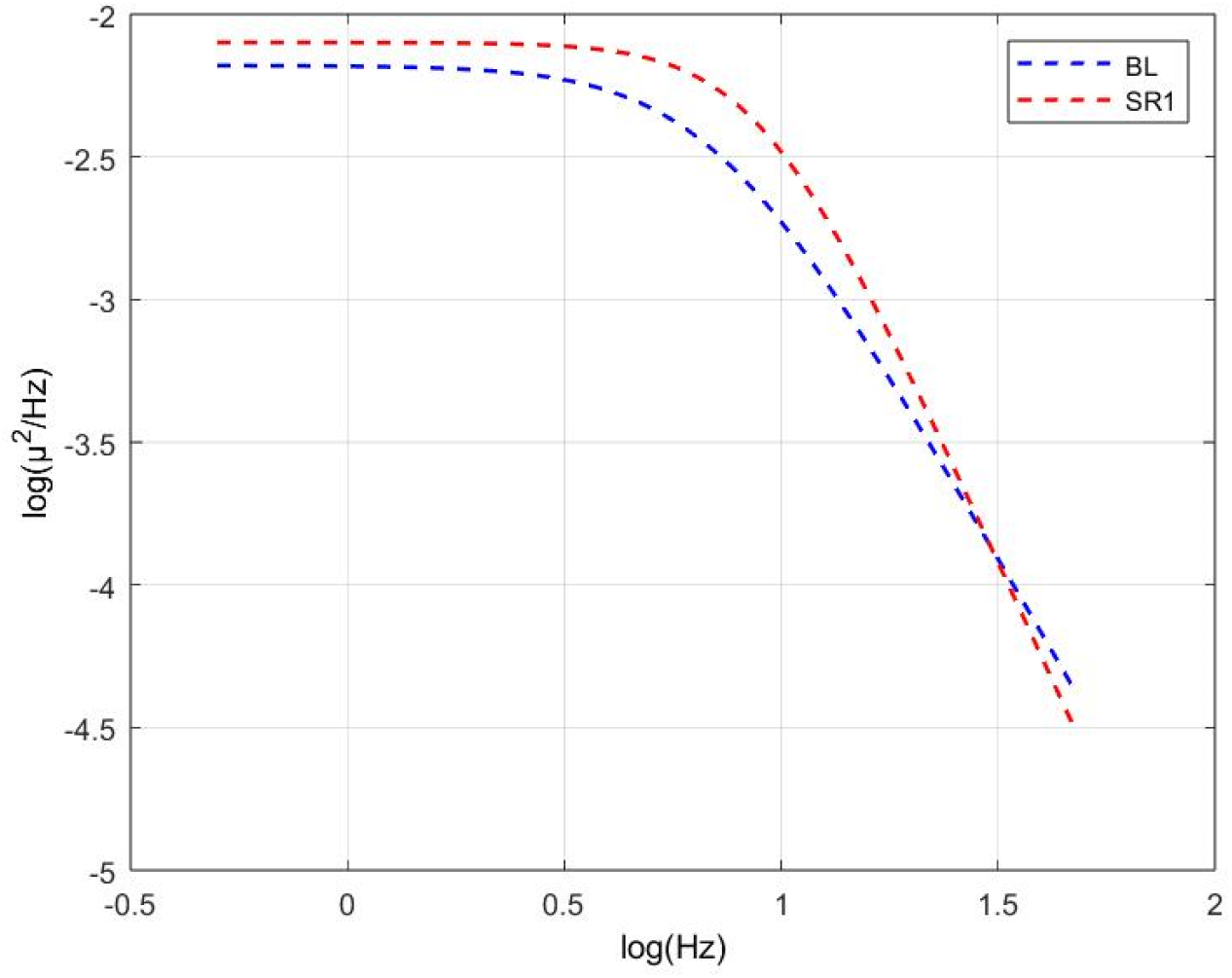
**Aperiodic fits of subject 1 over 19:00-20:00 BL and SR1 recordings**

**Figure 7.**
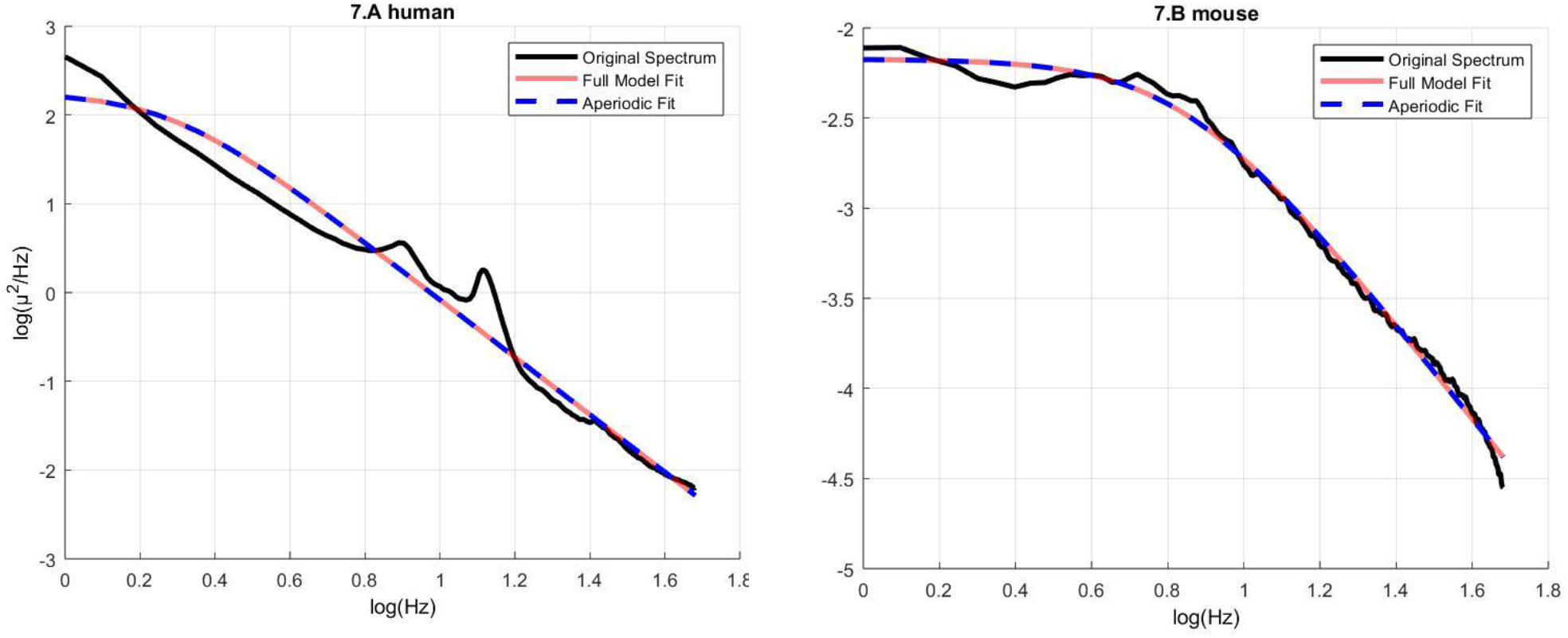
**The depiction of the comparative and species-specific perspective of the spectral knee issue in surface electrophysiological signals. A. The log-log representation of the scalp-recorded EEG of a healthy human volunteer (age: 27 years, gender: female, recording location: C4-A1A2) and its parametrization performed by the FOOOF procedure. Note the lack of the spectral knee in the displayed frequency range (1-48 Hz). B. The log-log representation of the ECoG of a mouse (subject 1, Baseline sleep during 19-20 hours, recording location: CP6 - interparietal bone) and its’ parametrization performed by the FOOOF procedure. Note the prominent spectral knee around 9 Hz frequency**

**Table 5.** A Descriptive statistics of Spectral Knee in WAKE.

|  | 19:00-20:00 | 20:00-21:00 | 21:00-22:00 | 22:00-23:00 | 23:00-24:00 |
| --- | --- | --- | --- | --- | --- |
| **Baseline** | 6.8(2.12) | 6.76(2.11) | 6.65(1.58) | 6.81(1.63) | 6.39(1.19) |
| **Sleep Restriction 1** | 7.44(1.57) | 7.21(2.11) | 7.19(1.93) | 7.85(1.53) | 6.41(1.25) |
| **Sleep Restriction 3** | 6.97(1.99) | 6.96(2.32) | 6.97(1.61) | 6.99(2.14) | 6.48(1.41) |
| **Sleep Restriction 5** | 6.89(2.04) | 7.08(2.05) | 6.85(1.69) | 7.21(2.05) | 7.55(2.09) |
| **Sleep Recovery 1** | 6.58(2.43) | 6.62(2.47) | 6.28(2.21) | 7.11(2.01) | 6.85(2.25) |
| **Sleep Recovery 3** | 7.29(2.2) | 7.09(1.94) | 7.09(1.5) | 6.95(1.77) | 7.18(1.78) |

**Table 5.** B Descriptive statistics of Spectral Knee in NREM.

|  | 19:00-20:00 | 20:00-21:00 | 21:00-22:00 | 22:00-23:00 | 23:00-24:00 |
| --- | --- | --- | --- | --- | --- |
| **Baseline** | 9(0.61) | 9(0.7) | 9(0.45) | 8.93(0.39) | 9(0.4) |
| **Sleep Restriction 1** | 9.48(0.63) | 9.38(0.55) | 9.31(0.49) | 9.16(0.66) | 9.21(0.68) |
| **Sleep Restriction 3** | 9.16(0.78) | 9.33(0.86) | 9.19(0.91) | 8.98(1.05) | 9.16(1) |
| **Sleep Restriction 5** | 9.44(0.74) | 9.35(0.9) | 9.33(0.76) | 9.38(0.86) | 9.39(0.74) |
| **Sleep Recovery 1** | 8.74(1.02) | 8.8(1.02) | 8.83(0.77) | 8.94(0.85) | 9.08(0.89) |
| **Sleep Recovery 3** | 8.87(0.82) | 8.79(0.9) | 8.85(0.79) | 8.92(0.71) | 8.97(0.62) |

**Table 5.** C Descriptive statistics of Spectral Knee in REM.

|  | 19:00-20:00 | 20:00-21:00 | 21:00-22:00 | 22:00-23:00 | 23:00-24:00 |
| --- | --- | --- | --- | --- | --- |
| **Baseline** | 9.89(1.34) | 10.09(1.1) | 10.17(1.19) | 9.79(1.11) | 9.94(0.94) |
| **Sleep Restriction 1** | 10.2(1.75) | 10.65(1.53) | 9.98(0.64) | 10.41(1.43) | 10.06(1.19) |
| **Sleep Restriction 3** | 10.85(1.74) | 10.39(1.43) | 10.61(1.27) | 10.37(1.44) | 10.52(1.33) |
| **Sleep Restriction 5** | 11.06(1.52) | 10.89(1.5) | 10.61(1.34) | 10.19(1.15) | 10.04(0.95) |
| **Sleep Recovery 1** | 9.46(1.78) | 9.57(1.3) | 9.97(1.39) | 10.03(1.47) | 9.58(1.19) |
| **Sleep Recovery 3** | 9.92(1.35) | 10.19(1.02) | 10.07(1.11) | 10.12(1.12) | 9.8(1.47) |

**Table 5.** D Descriptive statistics of Spectral Knee in ALL recorded periods.

|  | 19:00-20:00 | 20:00-21:00 | 21:00-22:00 | 22:00-23:00 | 23:00-24:00 |
| --- | --- | --- | --- | --- | --- |
| **Baseline** | 8.33(0.97) | 8.29(0.87) | 8.31(0.64) | 8.45(0.53) | 8.24(0.5) |
| **Sleep Restriction 1** | 8.98(0.78) | 8.65(0.93) | 8.58(0.86) | 8.86(0.87) | 8.14(0.62) |
| **Sleep Restriction 3** | 8.23(1.37) | 8.36(1.25) | 8.47(1.11) | 8.43(1.3) | 8.07(1.08) |
| **Sleep Restriction 5** | 8.13(1.61) | 8.32(1.61) | 8.3(1.19) | 8.63(1.23) | 8.68(1.04) |
| **Sleep Recovery 1** | 7.9(1.44) | 7.8(1.39) | 7.64(1.3) | 8.33(1.14) | 8.1(1.3) |
| **Sleep Recovery 3** | 8.25(1.26) | 8.11(1.3) | 8.14(0.88) | 8.39(0.94) | 8.41(0.88) |

note: Table 5.A, 2.B, 2.C and 2.D contains the descriptive statistics of Spectral Knee in WAKE, NREM, REM episodes, and when all scores were considered (ALL), in the following format: MEAN (STANDARD DEVIATION).

The spectral knee for WAKE (Table 5.A) and aggregated segments (Table 5.D; unifying WAKE, NREM and REM) did not display significant main effects. However, hypothesis-testing of REM segments (Table 5.C, Figure 8.) revealed significant main effect of HOURS (F(4, 16) = 11.755; p < .001; η_p_^2^ = 0.75), which was further dissected by post-hoc tests showing that in terms of the levels of the HOURS repeated measure factor the following contrasts were significant or marginally significant: 19-20 vs.

**Figure 8.**
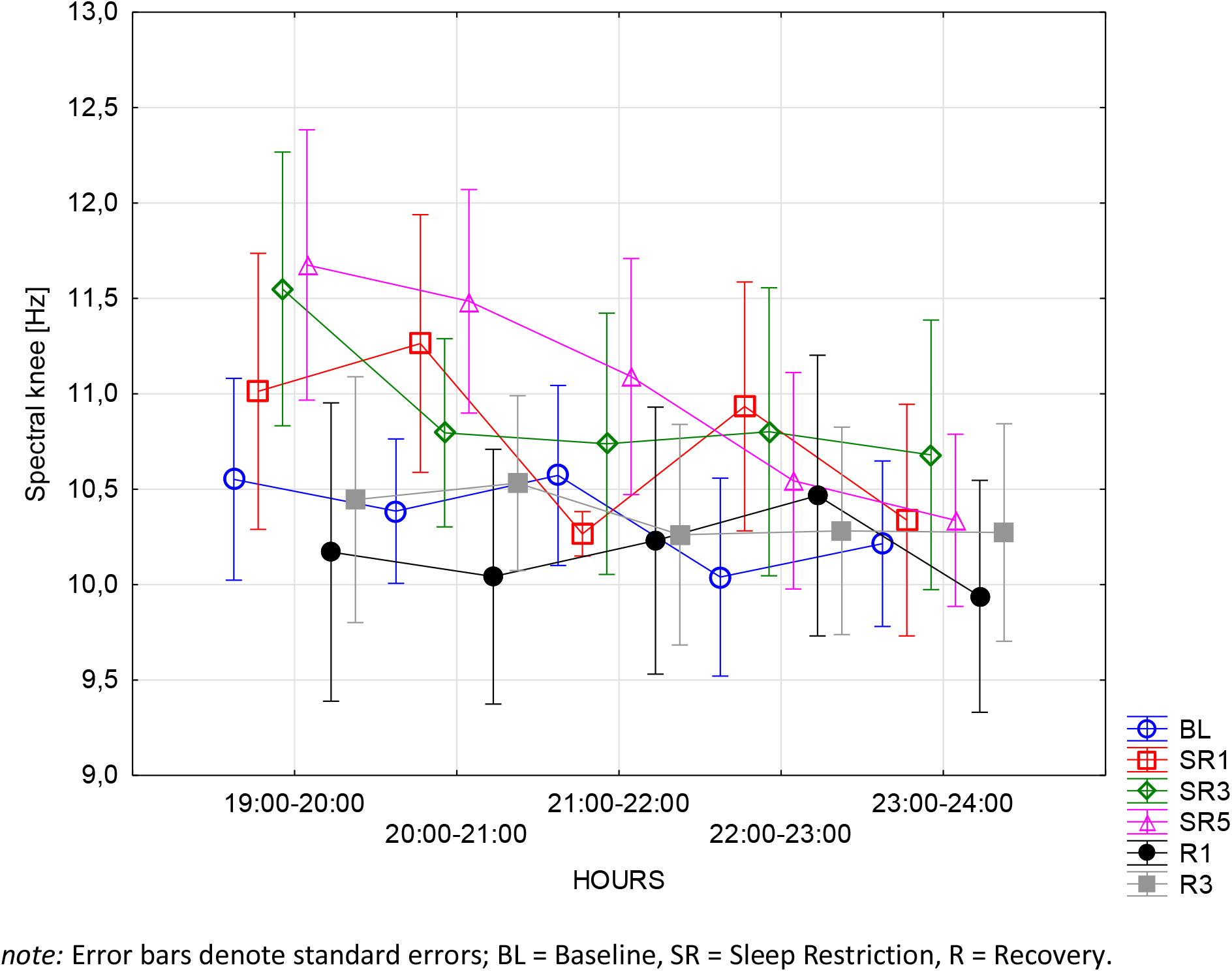
Spectral knees according to the FOOOF procedure as measured in ECoG recordings performed during REM periods of sleep.

21-22 (t = 3.8857; p = .0131; d = 0.2769), 22-23 (t = 4.0441; p = .0094; d = 0.2882), 23-24 (t = 6.2675; p < .001; d = 0.4466), and 20-21 vs 23-24 (t = 4.7072; p = .0023; d = 0.3354). However, the interaction between the independent variables was not found to be significant. Lastly, for NREM the spectral knee displayed no significant main effects.

### Normalized Spectral Entropy

No significant nor marginally significant DAY and HOUR effects in terms of spectral entropy were revealed for WAKE, NREM, REM, or aggregated segments.

**Table 6.**
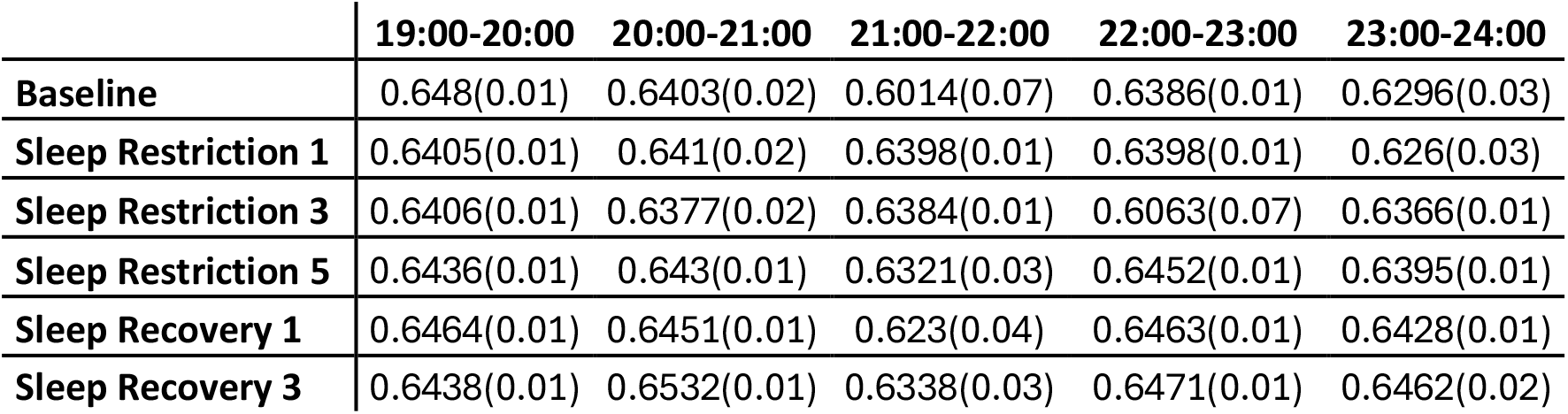
A Descriptive statistics of Normalized Spectral Entropy in WAKE.

|  | 19:00-20:00 | 20:00-21:00 | 21:00-22:00 | 22:00-23:00 | 23:00-24:00 |
| --- | --- | --- | --- | --- | --- |
| **Baseline** | 0.648(0.01) | 0.6403(0.02) | 0.6014(0.07) | 0.6386(0.01) | 0.6296(0.03) |
| **Sleep Restriction 1** | 0.6405(0.01) | 0.641(0.02) | 0.6398(0.01) | 0.6398(0.01) | 0.626(0.03) |
| **Sleep Restriction 3** | 0.6406(0.01) | 0.6377(0.02) | 0.6384(0.01) | 0.6063(0.07) | 0.6366(0.01) |
| **Sleep Restriction 5** | 0.6436(0.01) | 0.643(0.01) | 0.6321(0.03) | 0.6452(0.01) | 0.6395(0.01) |
| **Sleep Recovery 1** | 0.6464(0.01) | 0.6451(0.01) | 0.623(0.04) | 0.6463(0.01) | 0.6428(0.01) |
| **Sleep Recovery 3** | 0.6438(0.01) | 0.6532(0.01) | 0.6338(0.03) | 0.6471(0.01) | 0.6462(0.02) |

**Table 6.**
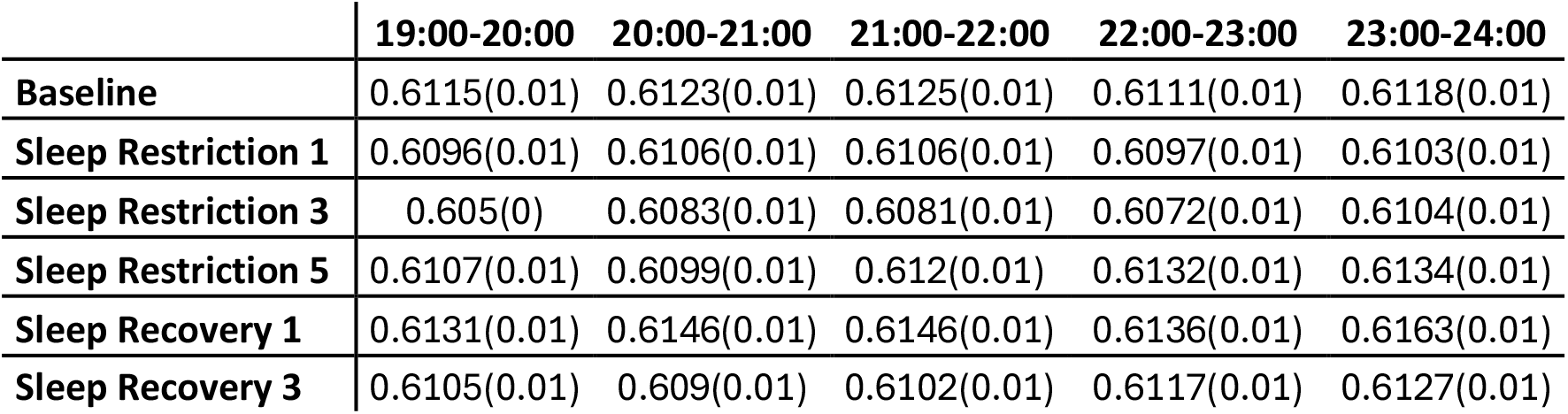
B Descriptive statistics of Normalized Spectral Entropy in NREM.

|  | 19:00-20:00 | 20:00-21:00 | 21:00-22:00 | 22:00-23:00 | 23:00-24:00 |
| --- | --- | --- | --- | --- | --- |
| **Baseline** | 0.6115(0.01) | 0.6123(0.01) | 0.6125(0.01) | 0.6111(0.01) | 0.6118(0.01) |
| **Sleep Restriction 1** | 0.6096(0.01) | 0.6106(0.01) | 0.6106(0.01) | 0.6097(0.01) | 0.6103(0.01) |
| **Sleep Restriction 3** | 0.605(0) | 0.6083(0.01) | 0.6081(0.01) | 0.6072(0.01) | 0.6104(0.01) |
| **Sleep Restriction 5** | 0.6107(0.01) | 0.6099(0.01) | 0.612(0.01) | 0.6132(0.01) | 0.6134(0.01) |
| **Sleep Recovery 1** | 0.6131(0.01) | 0.6146(0.01) | 0.6146(0.01) | 0.6136(0.01) | 0.6163(0.01) |
| **Sleep Recovery 3** | 0.6105(0.01) | 0.609(0.01) | 0.6102(0.01) | 0.6117(0.01) | 0.6127(0.01) |

**Table 6.**
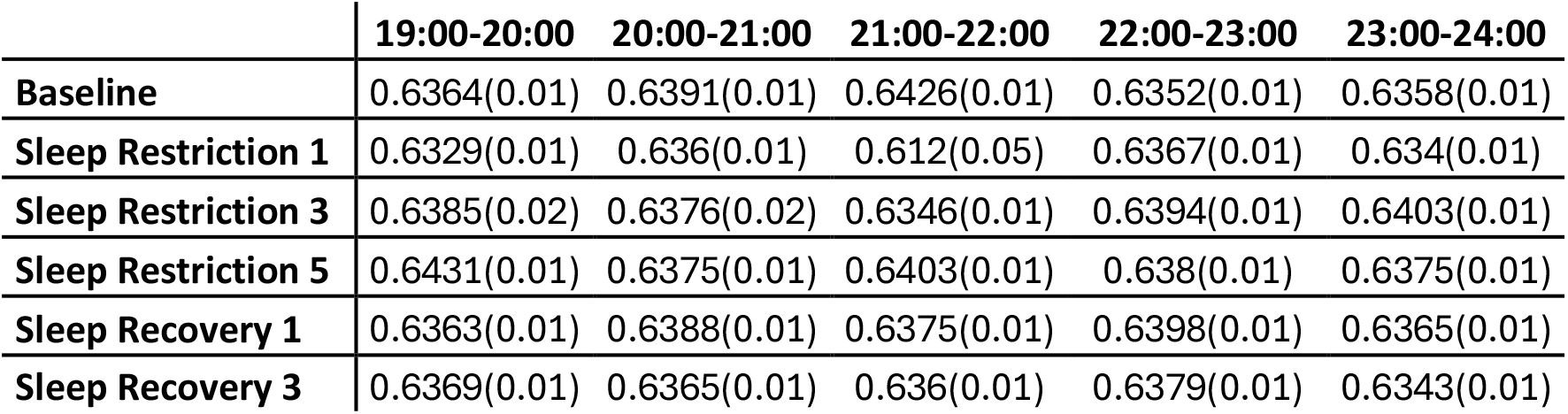
C Descriptive statistics of Normalized Spectral Entropy in REM.

|  | 19:00-20:00 | 20:00-21:00 | 21:00-22:00 | 22:00-23:00 | 23:00-24:00 |
| --- | --- | --- | --- | --- | --- |
| **Baseline** | 0.6364(0.01) | 0.6391(0.01) | 0.6426(0.01) | 0.6352(0.01) | 0.6358(0.01) |
| **Sleep Restriction 1** | 0.6329(0.01) | 0.636(0.01) | 0.612(0.05) | 0.6367(0.01) | 0.634(0.01) |
| **Sleep Restriction 3** | 0.6385(0.02) | 0.6376(0.02) | 0.6346(0.01) | 0.6394(0.01) | 0.6403(0.01) |
| **Sleep Restriction 5** | 0.6431(0.01) | 0.6375(0.01) | 0.6403(0.01) | 0.638(0.01) | 0.6375(0.01) |
| **Sleep Recovery 1** | 0.6363(0.01) | 0.6388(0.01) | 0.6375(0.01) | 0.6398(0.01) | 0.6365(0.01) |
| **Sleep Recovery 3** | 0.6369(0.01) | 0.6365(0.01) | 0.636(0.01) | 0.6379(0.01) | 0.6343(0.01) |

**Table 6.** D Descriptive statistics of Normalized Spectral Entropy in ALL recorded periods.

|  | 19:00-20:00 | 20:00-21:00 | 21:00-22:00 | 22:00-23:00 | 23:00-24:00 |
| --- | --- | --- | --- | --- | --- |
| **Baseline** | 0.6204(0.01) | 0.6209(0.01) | 0.616(0.01) | 0.6188(0.01) | 0.6179(0.01) |
| **Sleep Restriction 1** | 0.6178(0.01) | 0.6198(0.01) | 0.6214(0.01) | 0.6182(0.01) | 0.6171(0.01) |
| **Sleep Restriction 3** | 0.62(0.01) | 0.6209(0.01) | 0.6204(0.01) | 0.6119(0.01) | 0.6228(0.01) |
| **Sleep Restriction 5** | 0.6288(0.01) | 0.627(0.01) | 0.6248(0.01) | 0.6273(0.01) | 0.6236(0.01) |
| **Sleep Recovery 1** | 0.6238(0.01) | 0.6264(0.01) | 0.6237(0.01) | 0.6243(0.01) | 0.6288(0.01) |
| **Sleep Recovery 3** | 0.6199(0.01) | 0.6221(0.01) | 0.6212(0.01) | 0.6203(0.01) | 0.6227(0.01) |

note: Table 3.A, 2.B, 2.C and 2.D contains the descriptive statistics of Spectral Intercept in WAKE, NREM, REM episodes, and when all scores were considered (ALL), in the following format: MEAN (STANDARD DEVIATION).

## Discussion

### Spectral Slope, Spectral Intercept & Normalized Spectral Entropy

The present study investigates the sleep deprivation-related neurophysiological signals (electrocorticography, ECoG) of homeostatic adjustments during sleep, using the classical and a novel, data-driven signal processing method, and an information theoretical approach. The spectral slope (the steepness of the log-log power spectrum) compared to SWA (slow wave activity; 0.75 – 4.5 Hz), better captures the effects of prolonged wakefulness on sleep homeostasis. More precisely, suggesting the robustness of the spectral slope, it captures the increase of sleep deprivation-related enhanced sleep depth during both REM and NREM phases, as opposed to SWA only showing sleep homeostatic adjustments during NREM phase; in addition, the spectral slope reflects on sleep homeostasis outside the frequency range of what is traditionally considered a biomarker of sleep depth. In other words, the spectral slope appears to be a more composite index of sleep depth than the SWA. In addition, the spectral intercept, that is the point at which the model touches the ordinate-axis on the log-log transformed plane, reflects on sleep deprivation-related homeostatic processes during REM and NREM, just as the spectral slope. This is easily justified by the correlation between the spectral slope and the spectral intercept (16). Finally, the results suggest that normalized spectral entropy fails to differentiate between baseline sleep and sleep following sleep deprivation in mice.

The interpretation of the spectral slope is, however, a non-trivial matter. The surface neurophysiological power spectrum follows a tail-heavy exponential distribution, which upon a double logarithmic transformation linearizes, enabling the fit of a first-degree polynomial. As the domain is the log frequency bins, the slope of the linear curve can be considered reflecting on the ratio of slow oscillations to fast oscillations; in which case a steeper curve, that is higher absolute values of the spectral slope, means that low-frequency oscillations are relatively more dominant compared to high-frequency oscillations. On the other hand, a flat curve reveals the enhancement of fast oscillations. Therefore, in the context of sleep homeostasis, the steepness of the curve fitted over the power spectrum is, at heart, an expression of sleep depth (16). The Fractal and Oscillatory Adjustment Model (23) (FOAM) is a theoretical framework (a response to the scientific results aggregating since the conception of the two-process model) integrating homeostatic sleep pressure, circadian rhythms, and ultradian cycles into a comprehensive model. That is to say, it integrates the concept of fractal-like patterns with oscillatory processes to explain how the regulation of the sleep- wake cycle is achieved. In the context of FOAM, the spectral slope is a key parameter characterizing the scaling properties of the brain. Specifically, it reflects the fractal nature of neural background activity and sleep homeostasis. Therefore, the spectral slope may change in response to sleep pressure, making it a marker of sleep-wake history.

Furthermore, the spectral slope can be understood in terms of critical behaviour. Criticality, in essence, is a macroscopic feature that is the product of a balance between micro-level processes of the states in which a system can exist. Said balance corresponds to the phase transition between these states, and therefore, criticality is a characteristic of phase transitions, at which new properties emerge (7,54,55) – in terms of the brain, these properties manifest themselves in optimal computing, such as long-range communication or maximal information transfer between neuronal populations. Systems expressing scale-invariant phenomena may be characterized by criticality (8). Thus, the power-law scaling exponent (spectral slope) indexing fractal behaviour can be considered an index of critical dynamics. Investigating criticality involves the identification of an order and a control parameter. The order parameter is the macroscopic property characteristic of a given phase, while changes of the control parameter drive the fluctuations of the order parameter (7). In the context of the present study, **the order parameter is the spectral slope**, and **the control parameter is the experimental manipulation** (baseline sleep vs. sleep following prolonged wakefulness). Wakefulness is speculated to drive the brain towards supercritical behaviour, resulting in continuously flattening power spectra (1). Therefore, one may infer that sleep deprivation, that is inappropriately prolonged wakefulness, should exacerbate this phenomenon. The organic consequence of such is elevated sleep pressure resulting in a need for compensation, and thus a more pronounced movement towards subcriticality should be apparent in order for the system (the brain) to self-organize back into criticality. In other words, recovery sleep followed by sleep deprivation should bear signatures of pronounced subcriticality, such as a steeper power spectrum in the log-log space.

From an oscillatory point of view, supercriticality corresponds to a flat power spectrum, also called white noise, while subcriticality to a 1/f^2^ power spectrum, also called brown noise (56). Curiously, in between the flat white noise and the fast decaying brown noise lies the hyperbolic function in the form of 1/f, also called pink noise, which happens to be a signature fractal process of complex systems maintaining criticality possibly by a mechanism denoted self-organized criticality (57). Therefore, in accordance with our results, the steep power spectra characteristic to sleep following prolonged wakefulness may index a compensatory shift towards subcritical behaviour aimed to overcome the force exerted by sleep deprivation, which drove the brain towards supercriticality. However, the scaling exponents (spectral slopes) we found (Table 4.A-D) are pronouncedly outside of the range one would expect. Namely, the power spectrum of a system existing at criticality should be characterized by a scaling exponent approximating -2. This discrepancy, however, may be justified by the shift of the spectral knee towards higher frequencies (∼9 Hz; see below) rendering the scale-free portion of the power spectrum to decay faster and resulting in scaling exponents below the expected value. In addition, the brain’s tendency to organize into a subcritical regime and the relative simplicity of the rodent brain may also serve as arguments in favour of presenting our results in terms of the critical brain hypothesis. It has been previously suggested that the brain operates at a slightly subcritical regime during normal wake activity (1,25), also supported by human intracranial recordings of neuronal avalanches (58). Existing in a somewhat subcritical state allows the brain to avoid excessive noise in neural networks. Although near criticality offers maximal flexibility and responsiveness, a slightly subcritical regime provides a balance between stability and flexibility, minimizing the risk of runaway excitations or chaotic behaviour, and ensuring that neural responses are more stable and predictable. Moreover, the complexity of neural dynamics plays a significant role in whether the brain can operate near criticality or whether it is constrained to function in a suboptimal state. Rodent brains have simpler neural architecture, with fewer neurons and less intricate connectivity patterns as compared to human brains (59,60). The reduced complexity may limit the ability of rodent brains to exhibit the neural coordination necessary for criticality. In addition, the cognitive demands on a rodent are generally less diverse and complex than those on a human. Rodent brains are optimized for survival tasks like navigating, foraging, and escaping predators. These tasks require more ordered, predictable, stable neural dynamics that favor subcritical states – operating in a subcritical regime helps minimize metabolic costs. While energy efficiency is important in the human brain, its larger metabolic budget allows for more neural variability and activity. This increased metabolic capacity may support long-range synchrony, information integration, and flexibility, allowing the brain to explore a wider range of states. Thus, it can afford the energy costs of operating closer to criticality while still balancing metabolic demands.

### Spectral Knee

In light of the considerable shift towards higher frequencies of the spectral knee (∼ 9-10 Hz), the performance of the spectral slope to capture sleep homeostasis is noteworthy, as the fitted curve does not reflect on the frequency ranges (SWA: 0.75 – 4.5 Hz or delta power: 1 – 4 Hz), generally considered to be a biomarker of sleep homeostasis. To our knowledge, this right shift has only been reported in rats by Zhang et al. (61), however, there is no further discussion of the topic. Power law relationships in biological sciences concerned with scaling are rather pervasive (62) and are generally referred to as allometric equations (63,64). It is our suspicion that the right shift of the coloured- noise defined segment of the power spectrum is allometric in nature. In other words, the right shift in mice, compared to humans, is due to their smaller brain sizes. Mathematically this can be formalized such that

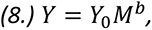

where Y is a biological function (e.x. spectral knee), Y_0_ is a constant, M is body size, and b is the allometric exponent; these parameters are easily obtained by a linear fit

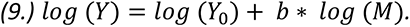

The power-law based allometric scaling can be investigated in terms of anatomic features and homeostatic processes as well (62). In the case of sleep homeostasis indexed by surface neurophysiological signals, the above equation may translate to plotting the spectral knee against body size. That is to say, with smaller body sizes come smaller brains, which means there is less matter that becomes synchronized, disproportionately affecting slow oscillations, resulting in flat power spectra in the slow frequency range. Given the tendency of biological systems to organize hierarchically and as a result to express fractal scaling (65,66), the spectral knee may signify the maximum capacity at which a brain is capable of producing (or required to produce) slow oscillations enabling long-range communication of neuronal populations (67) – another sign of criticality (7,68). This coheres with the theoretical consideration summarized by the so called Matérn process (69,70). The Matérn process is a generalization of the Gaussian process, with a covariance function defined by the Matérn kernel. Unlike fractional Brownian motion, which models power-law behavior, the Matérn process captures a low-frequency plateau, reflecting large-scale energy contributions (71). Therefore, the frequency domain properties of the Matérn process make it a more accurate method for turbulent modeling, as certain physical systems tend to generate a plateau around the lower frequencies, producing a kind of broken power-law with white and coloured noise in the low and the high frequency regions of the spectrum, respectively (71). In essence, the point at which the power spectrum breaks and starts to exhibit a decaying pattern, describes the physical limits of the system (71).

## Conclusion

We found that the fractal parameters (spectral slope & spectral intercept) of the neurophysiological power spectrum have better performance in capturing sleep deprivation-related homeostatic adjustments compared to the classical SWA approach. This is apparent as the fractal parameters reflect on sleep homeostasis irrespective of sleep staging, that is, sleep deprivation is indexed during both REM and NREM. Moreover, the shift of the spectral knee towards higher frequencies (∼ 9-10 Hz) renders the spectral slope unreflective of SWA, which is rather intriguing considering the history of SWA in the study of sleep homeostasis.

## Acknowledgments

Research supported by the Ministry of Culture and Innovation in Hungary (TKP2021-EGA-25). Research supported by the Ministry of Culture and Innovation in Hungary (TKP2021-NKTA-47).

## Notes

### Competing Interest Statement

The authors have declared no competing interest.

